# Specialized Pericyte Subtypes in the Pulmonary Capillary

**DOI:** 10.1101/2024.05.09.593174

**Authors:** Timothy Klouda, Yunhye Kim, Seung-Han Baek, Mantu Bhaumik, Yu Liu, Tiffany Liu, Jianwen Que, Joseph C Wu, Benjamin A Raby, Vinicio de Jesus Perez, Ke Yuan

## Abstract

Pericytes (PCs) play crucial roles in capillary maturation, stability, and homeostasis. Impaired PC coverage and function are implicated in various diseases, including pulmonary arterial hypertension (PAH). Challenges investigating PC biology are largely due to the lack of a concise marker, resulting in difficulty distinguishing PCs from other mural cell populations, including smooth muscle cells (SMCs) and fibroblasts (FBs). Utilizing bioinformatic analysis and RNAscope, we identified HIG hypoxia-inducible domain family member 1B (*Higd1b*) as a unique and conserved gene marker for PCs and generated a novel knockin mouse line, *Higd1b-CreERT2*, which precisely labels PCs in the lung and heart. Human lung single-cell RNAseq suggested the presence of two *HIGD1B*+ PC subtypes with different functions. By lineage tracing pulmonary *Higd1b+* cells exposed to hypoxia *in vivo*, we identified Type 1 PCs remained in the capillary network, while Type 2 PCs accumulated in the arterioles and coexpressed SMC markers and increased levels of Vimentin, associated with focal adhesion pathways. These results suggest that Type 1 PCs are specialized for supporting capillary EC homeostasis and quiescent, while Type 2 PCs are lineage active and located close to the border zone of the arterioles and capillaries, which may be motile and transition to SMC-like cells in hypoxia-induced pulmonary hypertension. The discovery of PC-type specialization in capillaries transforms our understanding of the structure, function and regulation of pulmonary capillary circulation and their contribution to vascular remodeling.

## Introduction

Pericytes (PCs) are mural cells that reside in the circulatory system’s microvasculature and maintain direct contact with capillary endothelial cells (ECs).^1^ Embedded within the basement membrane, they have critical roles in capillary homeostasis, angiogenesis, immune surveillance, and vessel maturation.^2–5^ The loss of PC function or coverage is associated with numerous diseases, including Alzheimer’s, diabetic retinopathy, cancer, and pulmonary arterial hypertension (PAH).^6–8^ Despite their involvement in pathological processes across multiple organ systems, PCs’ contribution to disease development is elusive in many circumstances. A significant limitation in studying PC biology arises from lacking a specific and unique cell marker, making it difficult to distinguish them from other mural cell populations under physiological and pathological conditions.^9^

Commonly used cell markers to identify PCs include Chondroitin Sulfate Proteoglycan 4 (*CSPG4,* aka Neuroglial Antigen NG2) and Platelet-Derived Growth Factor Receptor-β (*PDGFRb* aka CD140b).^10,11^ However, these markers lack specificity and uniqueness to PCs, as they are co-expressed in various cell types across multiple organ systems, including vascular smooth muscle cells (SMCs) expressing *Acta2* and *Tagln*, FBs expressing *Col1a1* and *Fbln1*, and oligodendrocytes expressing *Cspg4* and *Olig2.*^12–15^ To distinguish PCs from other mural cell populations, investigators rely on these cell markers combined with the PC’s distinct morphology (oval cell bodies with multiple, thin, prolonged and elongated branches/processes) and location in the circulatory system (abundantly encircling capillaries with a small population residing on the capillary-arterioles and capillary venule borders).^7,9,16^ Recent advances in single-cell RNA sequencing (scRNA-seq) have revealed organ-specific gene expression in PCs, reflecting their multifaceted functions throughout the body.^17^ Due to these limitations, there is a critical need to identify and validate a PC-specific cell marker.

PCs play a crucial role in supporting the circulatory system and exhibit multipotent stem cell-like properties.^18,19^ A notable histopathological feature of PAH is the excessive proliferation and accumulation of SMCs in the distal arterioles.^20^ Our group has demonstrated that in response to chronic hypoxia (Hx), NG2+ mural cells contribute to developing pulmonary hypertension (PH) and vascular remodeling by accumulating in the microvasculature and transitioning into SMC-like cells.^7^ PCs’ adaptability in response to Hx and organ injury positions them as promising targets for cell-directed therapies, not only in PAH but also in other disease processes.^21^ Through analysis of murine and human scRNA-seq databases, our group has previously identified HIG1 hypoxia-inducible domain family member 1B (*Higd1b*) as a gene exclusively expressed in *Cspg4+/Pdgfrβ+* mural cells in the heart and lungs.^17^

In this study, we utilize additional scRNA-seq databases and spatial transcriptomic analysis from human and murine lungs and hearts to identify *Higd1b* as a PC-specific gene marker. We employed the Cre-LoxP and CRISPR techniques to construct a novel, tamoxifen-inducible *Higd1b-CreERT2* mouse model. Validation with reporter lines (*R26-tdTomato* and *R26-mTmG*) confirmed Higd1b-Cre+ cells effectively labeled PCs in abundance in the lung and heart, and some in skeletal muscle, connective tissue, retina, and brain without labeling any other mural cells. Lastly, through lineage tracing studies, we demonstrated that two subtypes of PCs marked by *Higd1b-Cre* exist in the pulmonary circulation. Type 1 PCs are quiescent on capillaries, and Type 2 PCs exhibit multipotent properties and accumulate in the arterioles after exposure to Hx. These PCs likely undergo a transition into SMC-like cells via Vimentin activation, thereby contributing to vascular remodeling and the development of Hx-induced PH. This discovery positions *Higd1b* as a potential and unique cell marker for studying PCs in cardiorespiratory and vascular diseases. Additionally, the identification of PC subtypes in both humans and rodents will empower future investigators to delve into the dynamic role of PCs in disease development and pave the way for disease-modifying therapies targeting specific subgroups of PCs.

## Results

### Identification of *Higd1b* as a unique PC marker in human and murine lungs and hearts

Using murine and human scRNA-seq datasets from multiple tissue types, we identified several PC organ specific markers, including *Kcnk3* (lung), *Rgs4* (heart), and *Higd1*b (lung and heart), whose expression was restricted to annotated stringent PC clusters co-expressing both *Cspg4 and Pdgfrb.*^17^ Of these candidates, *Higd1b* was particularly interesting because its expression was conserved in both human lung and heart tissues. We speculated that *HIGD1B* would distinguish human and murine PCs from other mural cell populations in the cardiorespiratory system.

To further explore *HIGD1B* as a potential marker for PCs in lung and heart tissues, we examined the expression of *CSPG4*, *PDGFRB*, and *HIGD1B* in the human (Human Lung Cell Atlas Core) and murine (Tabula Muris Senis) single-cell datasets. The Human Lung Cell Atlas is a publicly accessible database comprising scRNA-seq on 584,944 human lung cells and circulating blood collected from 107 healthy individuals.^22^ The Tabula Muris Senis compendium (https://tabula-muris.ds.czbiohub.org/) is a publicly available single-cell transcriptomic atlas covering the lifespan of Mus musculus, including data from 23 tissues and organs. For cardiac tissues, the Human Heart dataset consists of scRNAseq on 486,134 human cells from all heart compartments and 14 individuals of both sexes, ranging in age from 40 to 75.^23^ The Mouse Heart dataset is from the C57BL/6 wildtype mouse strain, which consists of 25,436 cardiac cells.^24^ Utilizing the original UMAP coordinates and cell type annotations from each dataset, we mapped the annotated PCs onto the UMAP. We assessed their expression levels for known PC markers -*CSPG4*(*Cspg4*), *PDGFRB*(*Pdgfrb*), and *HIGD1B*(*Higb1b*)-in both human and murine scRNA-seq lung datasets (**Fig 1A**). The expression of *HIGD1B* was found exclusive to pulmonary PCs, while *PDGFRB* was expressed in PCs at equal levels and was also present in multiple FB subtypes, indicating broader cellular expression. Although *CSPG4* was also exclusively expressed in PCs, less than 30% of annotated PCs expressed *CSPG4*, and its relative expression to *HIGD1B* was reduced by about half (**Fig 1B**). A similar trend regarding *Higd1b* expression was seen in the Tanula Muris Senis dataset, as *Higd1b* was exclusively expressed in PCs without labeling other mural cells, while *Pdgfrb* was highly expressed in the PC population but also more broadly in other mural cells and *Cspg4* was exclusive to PCs but at significantly lower levels (**Fig 1C**). The expression of *CSPG4(Cspg4)*, *PDGFRB(Pdgfrb)*, and *HIGD1B(Higb1b)* for all 20 annotated human cell types and 30 murine cells can be seen in **Fig S1A.** In human and mouse heart scRNAseq, all three genes-*CSPG4*, *PDGFRB*, and *HIGD1B*-demonstrated relatively high expression within PC populations (**Fig S1B**). However, *HIGD1B* expression was not exclusive to PCs, as we observed a subset of SMCs also expressing it. Notably, *Higd1b*’s expression, in comparison to *Cspg4* and *Pdgfrb*, was both high and more specific to mouse heart PCs (**Fig S1C**).

**Figure 1.**
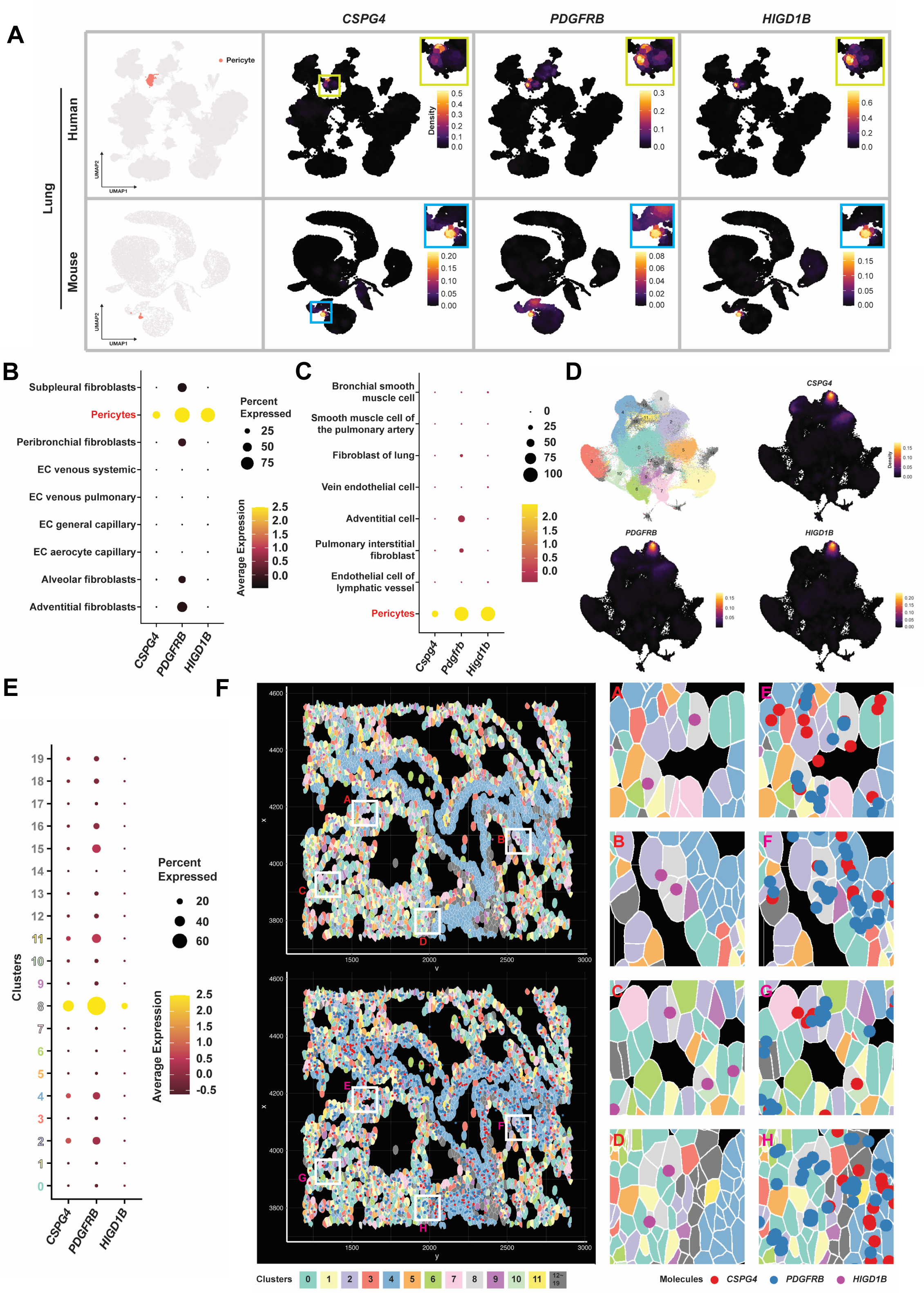
*HIGD1B*/*Higd1b* is identified as a unique and exclusive marker for human and murine PCs. (A) UMAP visualization of PC distributions within human and mouse lung tissues, using original UMAP coordinates and cell type annotations from the ‘Human Lung Cell Atlas (core)’ and ‘Tabula Muris Senis’. The three panels on the right reveal the expression of known PC markers CSPG4, PDGFRB, and HIGD1B, with a magnified image showing expression among annotated PCs in the top right corner. (B) Dot plot shows expressions of *CSPG4, PDGFRB,* and *HIGD1B* in nine different cell types from the Human Lung Atlas. Note the exclusive expression of *HIGD1B* in PCs compared to other mural and vascular cells. Comprehensive expression data across all annotated cell types are presented in Supplementary Figure 1. (C) Dot plot shows expressions of *Cspg4*, *Pdgfrb* and *Higd1b* in eight different vascular cell types from the Tabula Muris Senis compendium. As seen in the Human Lung Cell Atlas, the exclusive expression of *Higd1b* in PCs was found when compared to other mural and vascular cells. (D) UMAP shows 20 specific cell clusters derived from the spatial transcriptomic analysis using the ‘Xenium Human Lung Preview Data (Non-diseased Lung)’. PCs are annotated in Cluster 8 (light gray) based on expressions of *CSPG4*, *PDGFRB*, and *TRPC6*. The density plots specifically highlight the expression of *CSPG4*, *PDGFRB*, and *HIGD1B* across all clusters. (E) Dot plot illustrates the expressions of *CSPG4*, *PDGFRB*, and *HIGD1B* across 20 cell clusters identified by the spatial transcriptomic analysis of the ‘Xenium Human Lung Preview Data (Non-diseased Lung)’. Again, note the *HIGD1B* expression is exclusive to PCs (Cluster 8). (F) Detailed spatial plots for a selected lung tissue section (coordinates: x: 1200-2900, y: 3750-4550) utilizing ‘ImageDimPlot’ function in Seurat. This visualization focuses on well-defined stromal and capillary structures, showcasing the spatial distribution of cells in Cluster 8, predominantly consisting of PCs. Cells are color-coded by cluster, highlighting the localization of *CSPG4*(red), *PDGFRB*(blue), and *HIGD1B*(purple). Panels A-D show magnified lung microvasculature and the high expression of *HIGD1B* in PCs (light gray) annotated/colocalized in Cluster 8. The full lung tissue section can be found in Supplementary Figure 2.

To further explore the spatial distribution of PCs within lung tissue, we analyzed the spatial transcriptomic data from the ‘Xenium Human Lung Preview Data (Non-diseased Lung)’ provided by 10x Genomics. By projecting cells onto a two-dimensional UMAP space and performing unsupervised clustering, we identified 20 distinct cell clusters (**Fig 1D**). Among these, Cluster 8 exhibited high expression levels of *CSPG4*, *PDGFRB,* and *HIGD1B*. Notably, *CSPG4* and *HIGD1B* expression were specific to Cluster 8, whereas expression of *PDGFRB*, and to a lesser extent *CSPG4,* was observed in other clusters (**Fig 1E**). These findings were consistent with the expression patterns observed in the scRNA-seq datasets annotated with PCs, which suggested that Cluster 8 (light grey) predominantly consisted of PCs. To investigate the spatial arrangement and location of these cells within the lung tissue, we focused on a lung section with well-preserved architecture exhibiting clearly defined alveolar and capillary structures. Within this structure, we were able to label cell types based on prior cell clustering and identify the location of PCs within the lung (**Fig S2A&B**). After mapping lung cells to their respective clusters, we investigated a random area of the lung section (coordinates: x:1200-2900, y:3750-4550) to better visualize individual cell borders and localize the expression of *CSPG4* (red dots), *PDGFRB (*blue dots), and *HIGD1B (*purple dots) among all cell clusters captured within this region. *HIGD1B* expression showed a broader distribution and was predominantly localized in cells originating from Cluster 8 (**Fig 1F, Panel A-D)**. In contrast, *PDGFRB* was mingled with *CSPG4,* and both were predominantly around the smooth muscle layer of an artery when tracing the distribution pattern of Cluster 4(light blue) (**Fig 1F, Panel E-H**). In summary, using human and murine scRNA-seq datasets from lung and heart tissue along with spatial transcriptomics, we identified *HIGD1B* as a more specific marker for pulmonary PCs compared to *CSPG4* or *PDGFRB*.

### Construction of *Higd1b-CreERT2* knockin mouse line

To validate *Higd1b* as a PC-specific cell marker, we designed *Higd1b*-mRNA probes and performed RNAscope on *Cspg4-CreERTM::R26-tdTomato* (NG2-tdT) lungs. In precision cut lung slices(PCLSs), parenchymal tdT+ cells clearly demonstrated classical PC morphology, including an oval cell body and long punctate processes partially wrapping capillary ECs (**Fig 2A**). We applied *Higd1*b probes on OCT embedded NG2-tdT lungs and found 100% co-localization of *Higd1b* mRNA with endogenous tdT (**Fig 2B**). *Higd1b* positive punctate dots also stained some cells without tdT, suggesting incomplete tamoxifen induction of NG2-tdT recombination. In larger-sized arteries, no *Higd1b* staining was found in the smooth muscle layers (**Fig S3**). After RNAscope validation, we sought to generate a novel, tamoxifen-inducible, knockin mouse line using the Cre-LoxP system using CRISPR **(Fig 2C)**. A total of 47 pups were screened by PCR using Cre-specific primers, and two founders were identified, which were #12 and #25. The genotyping results shown were from mouse #25. To confirm target specific intact integration of the donor construct in the Higdb1 locus, for LHA the forward primer LF is designed in the intronic region between exon-1 and exon-2 and C1 reverse primer in 5’ region of Cre (1.9kb); the RHA is screened using forward primer C4 located in 3’ of Cre and reverse primer RR is outside of RHA in intron (2.3kb) (**Fig 2C**). Sanger sequencing of PCR products from left and right homology arms confirmed knock-in of *Higdb1-P2A-CRE-ERT2* construct in exon 4 (**Supple Table 1**).

**Figure 2:**
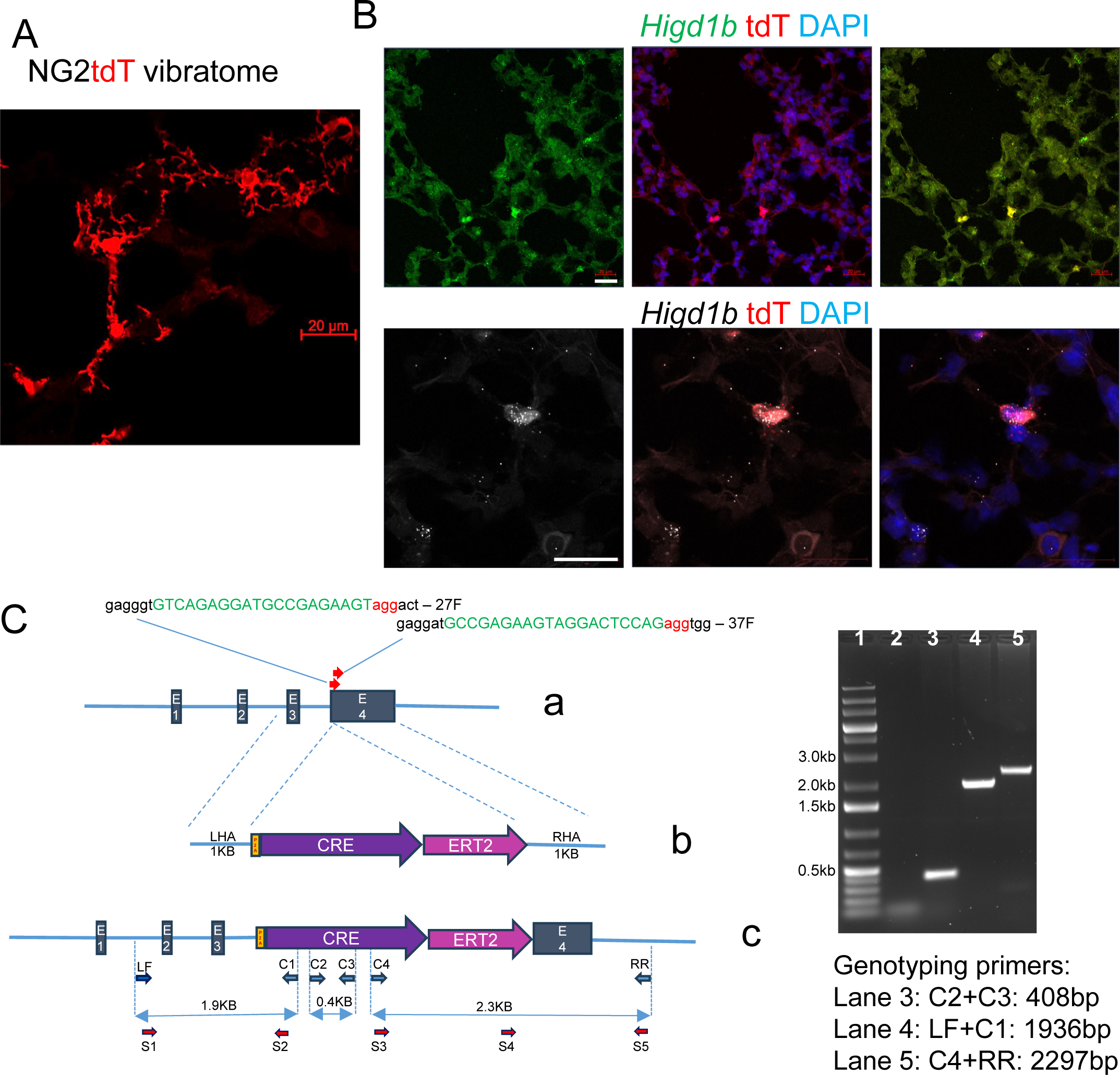
*Higd1b-CreERT2* knockin mouse line is constructed by CRISPR. (A) Precision cut lung slice(PCLSs) were obtained from *Cspg4-CreERTM::Ai14* (NG2-tdT) mice. Scale bar: 20µm. (B) RNAscope of the *NG2-tdT+/-* mouse OCT sections shows the coexpression of *Higd1b* mRNA (green and white) and the tdT reporter (red). Nuclei stained with DAPI (blue). Scale bar: 20 µm. (C) Schematic figure for Cre-ER insertion and PCR validation of insertion. a: *Higd1b* locus exon 1-4 and location of two gRNA and sequences in green, PAM sites in red and adjacent nucleotides in black after ATG. b: the Knock-in targeting construct: left (LHA) and right homology arms (RHA) flanked by P2A-CRE-ERT2, c. genomic structure after P2A-CRE-ERT2 knock-in. Primer pair LF+C1 identifies 5’ end, and C4+RR identifies ERT2 and 3’ end, C2+C3 Cre gene. Left: Two founder mice, #12 and #25, were identified by Cre-specific PCR using C2+C3 (lane 3) primers LF is located outside of LHA, C1 is located 5’ of Cre, C4 is located at the 3’ of Cre, and RR is outside of RHA.

### *Higd1b-Cre+* specifically labels PCs in the lungs and hearts *in vivo*

To further validate *Higd1b* sensitivity for PCs, we bred *Higd1b-CreERT2* with an *R26-tdTomato* (referred to as *Higd1b-tdT*) and an *R26-mTmG (*referred to as *Higd1b-mTmG*) reporter mouse and treated both with tamoxifen to confirm endogenous labeling of PCs.

Examination of PCLSs from *Higd1b-tdT* with confocal microscopy revealed endogenous reporter tdT cells with morphology representative of pulmonary PCs; a central, oval body with multiple, elongated processes (**Fig 3A**). These cells were found distributed through the capillaries of the pulmonary parenchyma but not in distal arterioles (>25 µm). More importantly, there was no co-expression of tdT positive (+) cells colocalized with either EC marker CD31 (green) or SMC marker SMA (white). **(Fig 3A, top and middle rows).** Furthermore, the majority of tdT+ cells were found to co-express PDGFRβ (green), although PDGFRβ staining was also seen in tdT negative(-) cells in arterioles/arteries, highlighting non-specificity for *Pdgfrb* to identify PCs **(Fig 3A, bottom row)**. We then evaluated precision cut heart slices from *Higd1b-tdT* due to our prior scRNA-seq results suggesting the expression of *Higd1b* in cardiac PCs^17^. We observed tdT+ cells stained for red-fluorescent protein (RFP) labeled PCs in capillaries throughout the myocardium, which did not coexpress SMA (**Fig 3B)**. Notably, IF staining and confocal microscopy from *Higd1b-mTmG* mice demonstrated similar findings, with cells endogenously labeled with green fluorescent protein (GFP) reporter in PC-shaped cells distributed throughout the parenchyma in direct contact with ECs (CD31). Compared to *Higd1b-tdT*, *Higd1b-mTmG* seemed to label more PC cytoplasmic processes. The majority of GFP-labeled cells coexpressed staining for PDGFRβ, but no visualized cells were positive for SMA **(Fig 3C).** Both *WT-tdT+/-* and *WT-mTmG+/-* mice injected with similar doses of tamoxifen and knockin mice (*Higd1b-td+/-* and *Higd1b-mTmG+/-)* without tamoxifen treatment did not express endogenous labeling for tdT or GFP **(Fig S4A&B),** suggesting recombined reporter color is driven by tamoxifen induction.

**Figure 3:**
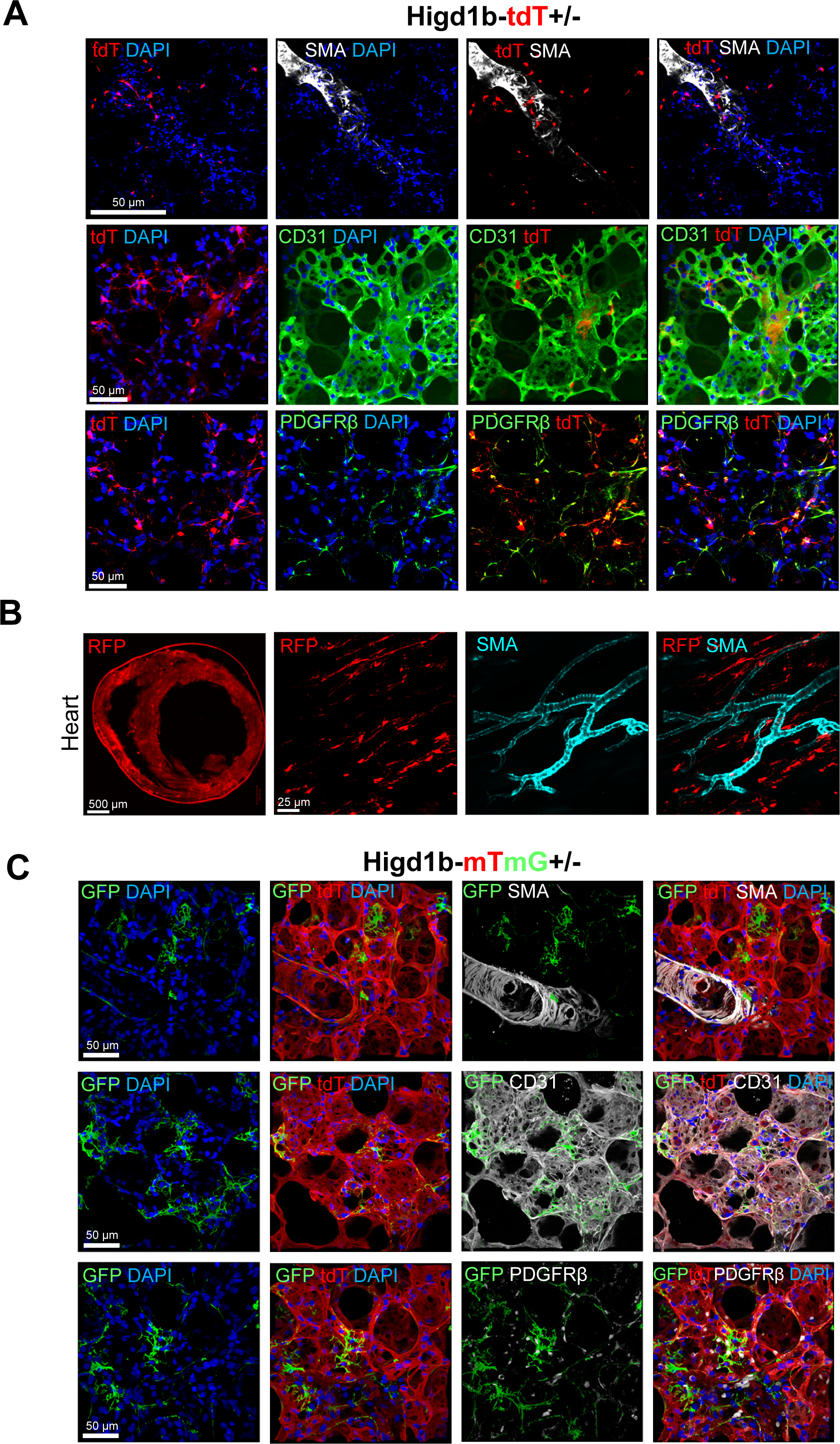
*Higd1b-Cre+* cells precisely label pulmonary and cardiac PCs. (A) Representative images of lung tissues from *Higd1b-tdT+/-* mice were stained for SMA (SMC marker: white), CD31 (EC marker: green), PDGFRβ (mural marker: green), and nuclei (DAPI: blue). PCs labeled with tdT reporter (red) are endogenous colors without antibody staining. Note the distinct morphology of tdT+ cells with an ovoid body and elongating processes wrapped around the capillary networks. tdT+ cells had no expressions of SMA or CD31. However, the majority (but not all) of tdT+ demonstrate coexpression with mural cell marker PDGFRβ. Scale bar: 50 µm. (B) Vibratome was prepared for heart tissue from *Higd1b-tdT+/-* mice after IF staining for RFP (left). Scale bar: 500 µm. The right panels show an area under increased magnification and RFP-positive cells without coexpression of SMA (cyan). Scale bar: 25 µm. (C) Representative images show GFP reporter labeled PCs (green) from lung tissue of *Higd1b-mTmG+/-* mice with staining for SMA (white, top panel), CD31 (white, middle panel), PDGFRβ (white, bottom panel), and DAPI (blue). Scale bar: 50 µm.

In addition to the cardiorespiratory system, PCs contribute to numerous diseases, including Alzheimer’s, diabetic retinopathy, renal fibrosis, and more in numerous organ systems^25^. We, therefore, examined additional tissues harvested from *Higd1b-tdT* mice for the endogenous labeling of PCs. Similar to heart and lung samples, confocal microscopy of skeletal muscle and connective tissues of descending aorta from *Higd1b-tdT+/-* mice revealed SMA-negative, tdT-positive PCs in direct contact with the microvasculature (**Fig S5A&B**) Retina from *Higd1b-tdT*+/-mice also demonstrated minimal labeling of PCs(∼4.6%) (**Fig S5C).** Finally, considering the importance of PCs to the maintenance of the blood-brain barrier, we evaluated murine brain tissue.^26^ Sections of *Higd1b-tdT+/-* brain demonstrated few tdT+ cells lining the blood vessels (**Fig S5D**). There was no consistent identification or expression of tdT in cells from either the kidney or liver of *Higd1b-tdT* mice (**Fig S6**).

### PCs translocate dynamically in 3wk hypoxia induced PH and recovery models

Utilizing NG2-tdT, our group previously showed that NG2+ mural cells accumulate in the microvasculature and contribute to the development of Hx induced PH.^7^ Since NG2 is a shared cell marker expressed in multiple mural cell populations, particularly in SMCs, we can not conclude that PC’s direct contribution to arteriole muscularization and vascular remodeling. Thus, we performed lineage-tracing studies using *Higd1b-tdT+/-* and *Higd1b-mTmG+/-* mice to precisely describe the contribution of PCs to vascular remodeling across different periods of Hx.

After verifying the appropriate and selective labeling of PCs in *Higd1b-tdT+/-* mice after tamoxifen injection and allowing rest for up to 2 weeks, we exposed mice to Hx (1wk, 2wks, 3wks) and harvested lung tissues.^27,28^ Using confocal microscopy and IF staining, we described morphology, distribution, and expression of SMC cell markers in *Higd1b-tdT+/-* positive cells in response to Hx. After 1wk of Hx, endogenously labeled tdT+ cells were found in close proximity to the distal arterioles (<50 µm). A small subset of tdT+ cells were found directly in muscularized vessels expressing positive SMCs (SMA+) staining. Compared to normoxic samples, PCs in the distal arterioles demonstrated a spindle-shaped morphology that closely resembled SMCs with reduced process length compared to normoxic PCs. The majority of PCs, however, were still found within the lung parenchyma and in direct contact with capillary ECs (**Fig 4A, top panel**). After prolonged exposure to Hx for up to 3 weeks, the quantity of tdT+ cells with spindle-shaped(shorter processes) morphology in the remodeled arterioles was increased, suggesting PCs were in direct contact with the muscularized vasculature. Notably, many PCs remained in the lung parenchyma and in direct contact with capillary ECs with potentially longer and increased surface areas of processes (**Figure 4A, bottom panel).** Additional IF staining for the mature SMC marker SMMHC in mice exposed to Hx revealed coexpression of tdT+ PCs with SMMHC and SMA (**Fig S7A**).

**Figure 4:**
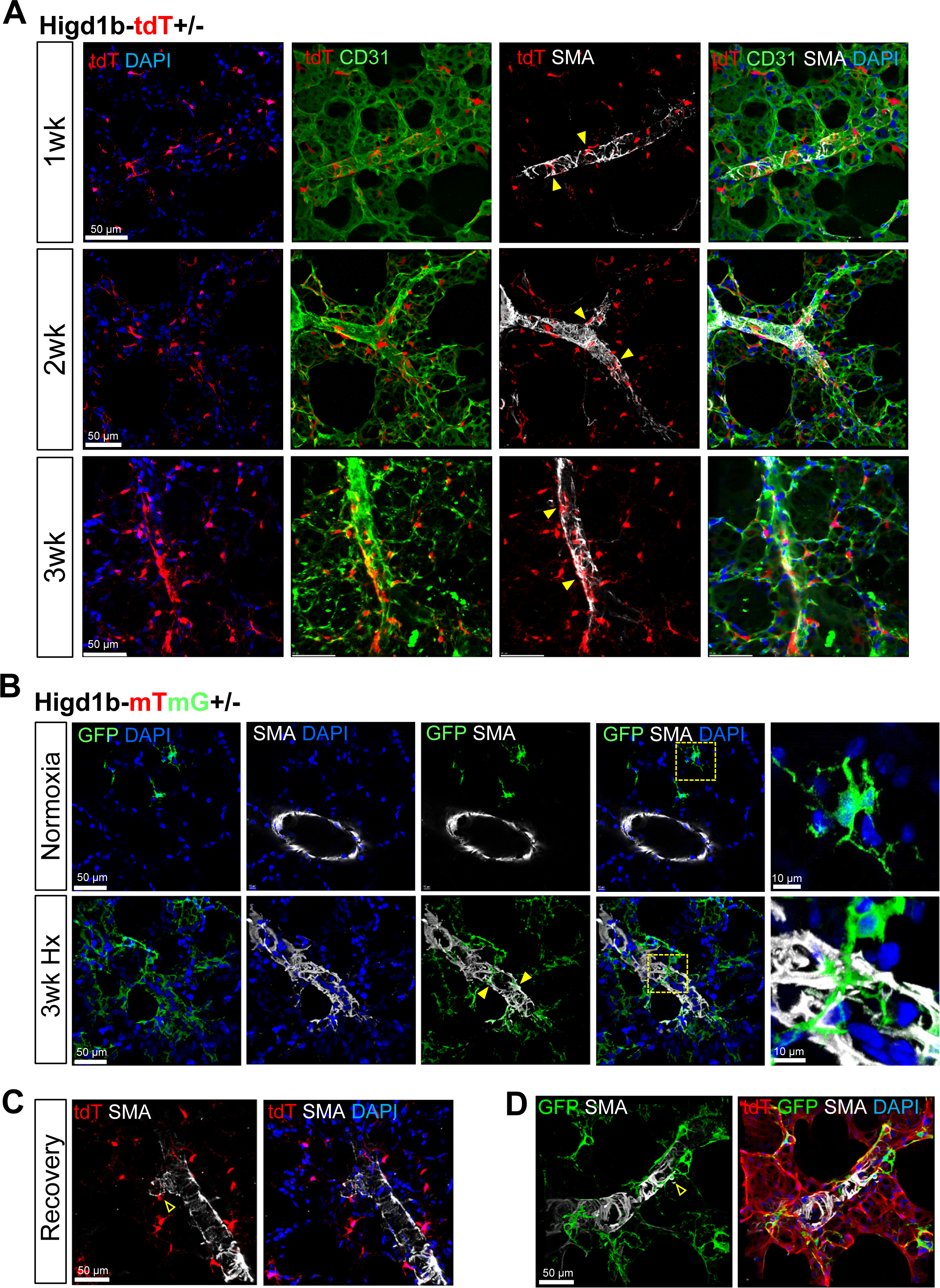
Lineage tracing shows that *Higd1b-Cre+* cells accumulate in muscularized distal arterioles at different hypoxic exposure times. (A) PCLSs from *Higd1b-tdT+/-* mice exposed to Hx stained for CD31 (EC marker: green), SMA (SMC marker, white), and nuclei (DAPI: blue). Note tdT+ (red) reporter cells without antibody labeling(indicated by yellow arrowheads) accumulated in the distal vasculature after 1 week of Hx and increased coverage on arterioles after prolonged exposure for up to 3 weeks. Scale bar: 50 µm. (B) Lineage tracing experiments for *Higd1b-mTmG+/-* mice with IF staining for SMA (white) and nuclei (DAPI). The yellow boxes demonstrate areas of increased magnification in the right panel, showing spindle-shaped morphology of GFP+ PCs (yellow arrowheads) coexpress SMA after exposure to 3wk of Hx in the distal vasculature compared to normoxic controls. Scale bar: 50 µm. (C) Lineage tracing for the recovery model using *Higd1b-tdT+/-* mice exposed to Hx then returned to normoxia for 3wks and stained for SMA (SMC marker, white), and nuclei (DAPI: blue). Note that the majority of tdT+ (red) reporter cells returned to the parenchymal capillary. Only a small number of tdT remained on the distal arterioles(hollow arrowheads). Scale bar: 50 µm. (D) Lineage tracing for the recovery model using *Higd1b-mTmG+/-* mice exposed to Hx then returned to normoxia for 3wks and stained for SMA (SMC marker, white), and nuclei (DAPI: blue). Note that the majority of GFP+ (green) reporter cells returned to the parenchymal capillary. Only a small number of GFP remained on the distal arterioles(hollow arrowheads). Scale bar: 50 µm.

Next, we performed similar lineage-tracing experiments in *Higd1b-mTmG+/-* mice to further validate our findings. In normoxic conditions, PCLSs from *Higd1b-mTmG+/-* mice demonstrated endogenously labeled GFP+ cells (PCs) within the lung parenchyma and morphology representative of a PC. No visualized GFP+ cells had coexpression for SMA or were located on arterioles (**Fig 4B, top panel**). After 3 wks of Hx, GFP+ was found accumulated in the distal arterioles (25-50 µm), expressing SMA and with a spindle-like shape compared to normoxic conditions (**Fig 4B, bottom panel**). The parenchymal GFP+ showed increased process coverage.

The chronic Hx mouse model is a well-established experimental PH model and all of the Hx-induced PH and pulmonary vascular alterations are fully reversed after 3 weeks of normoxia recovery (referred to as Recovery model).^29^ Upon returning hypoxic *Higd1b-tdT+/-* and *Higd1b-mTmG+/-* mice to normoxic conditions, we observed the loss of PC accumulation on distal arterioles and restorations of normal PC morphology similar to baseline conditions, suggesting a dynamic movement of PC in pathological conditions (**Fig 4C & D**). However, we noticed that a small portion (2.5%) of tdT+ or GFP+ cells still wrapped and resided in the very distal arterioles but no longer expressed SMA. We speculate that longer exposure to normoxia may result in a complete return of PC to their parenchymal location.

### Two PC subtypes are identified in capillary networks

A key finding in our fate-mapping experiments was the distinct location and morphological change of *Higd1b+* PCs after exposure to Hx. Since *Higd1b+* cells accumulated in remodeled arterioles or remained in the capillaries after exposure to Hx, we speculated that there may be different PC subtypes within the pulmonary capillaries. These subsets of PCs may have different roles in disease development and be distinguished from one another using known PC markers *Higd1b, Pdgfrb,* and *Cspg4*. Therefore, we sought to determine if such PC subsets could be identified, providing further insight into the role of PCs in the development of Hx-induced vascular remodeling using scRNA-seq and spatial transcriptomics.

To explore the heterogeneity within PC sub-populations in relation to *HIGD1B* expression, we first subset the annotated PC cluster from the “Human Lung Cell Atlas”. ^22^ Utilizing Harmony^30^ to remove the batch effect from sub-clustering, we identified four distinct PC sub-Clusters(**Fig 5A** and **Fig S8**). Notably, sub-Cluster 0 and 1 exhibited high expression levels of *HIGD1B*, whereas sub-Cluster 2 and 3 displayed relatively lower expression. Particularly, despite its lower *HIGD1B* expression, sub-Cluster 3 demonstrated a higher expression of *CSPG4* and *PDGFRB*.

**Figure 5:**
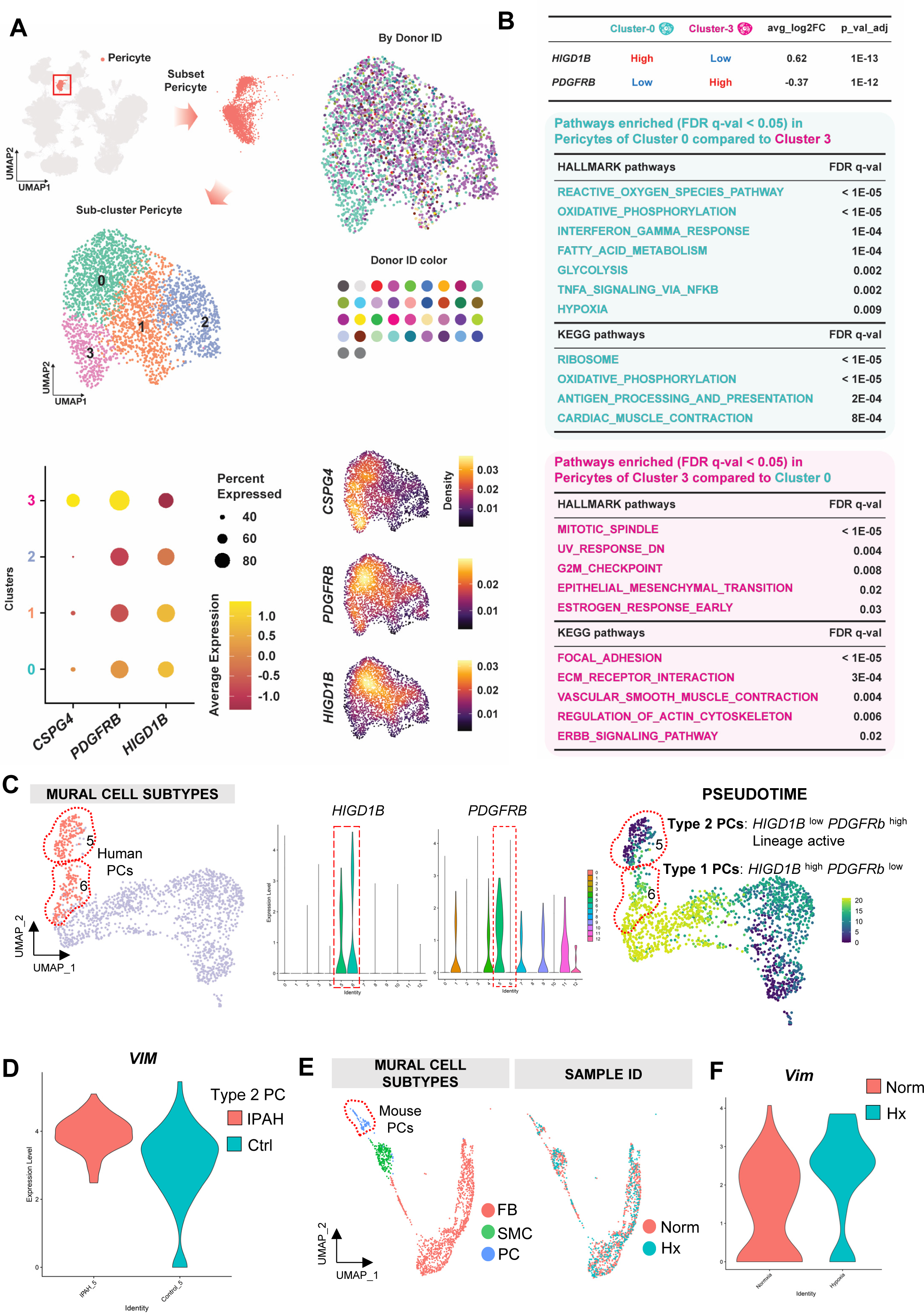
Two subtypes of PCs are identified using human lung scRNAseq. (A) UMAP visualization of four PC sub-populations within the annotated PC cluster from the ‘Human Lung Cell Atlas (core)’. The density and dot plots distinctly illustrate the expression levels of *CSPG4*, *PDGFRB*, and *HIGD1B*, revealing the heterogeneity within the PC sub-clusters based on these marker genes. (B) Gene Set Enrichment Analysis (GSEA) comparing PC sub-Cluster 0 and 3. Sub-Cluster 0, marked by significantly higher expression of *HIGD1B*, is characterized by enrichment in metabolic activity and stress response pathways. Sub-cluster 3, with a notably higher expression of *PDGFRB*, is associated with pathways involved in focal adhesion, cell cycle regulation and movement. A comprehensive list of enriched pathways with an FDR below 0.05 can be found in Supplementary Figure 9. (C) Human IPAH and control mural cell subtypes were re-clustered from the previously published analysis. Sub-clusters 5 and 6 were enriched with expression profiles of *HIGD1B* designated as pericyte clusters. The violin plot shows the gene expression level of *HIGD1B* in subclusters 5 and 6 but is higher in subcluster 6. The Violin plot shows the gene expression level of *PDGFRB* in sub-Clusters 5 and 6 but is higher in sub-Cluster 5. Pseudotime analysis shows Type 1 PCs (sub-Cluster 6, *HIGD1B^high^ PDGFRB^low^*) have a majority of late lineages (yellow), whereas Type 2 PCs (sub-Cluster 5, *HIGD1B^low^ PDGFRB^high^*) have a mixture of early (purple) and middle lineages (green). (C) The Violin plot shows the gene expression level of *VIMENTIN(VIM)* is increased in IPAH Type 2 PCs vs healthy donor Type 2 PCs in sub-Cluster 5. (E) The UMAP plot of mouse lung mural cells that underwent hypoxia vs. normoxia was reanalyzed from previously published work, and the PC cluster was identified in the dashed area. (F)The Violin plot shows the gene expression level of *Vimentin(Vim)* is increased in hypoxic PCs vs normoxic PCs.

To further dissect the molecular difference between PCs with high and low *HIGD1B* expression, we performed differential expression (DE) analysis between sub-clusters 0 and 3, utilizing the Wilcoxon rank-sum test. Subsequently, we conducted a pre-ranked Gene Set Enrichment Analysis (GSEA) to identify key pathways and genes distinguishing these sub-populations (**Fig 5B**). Our analysis revealed that the relative expression levels of *HIGD1B* were significantly higher in sub-Cluster 0, whereas the relative expression levels of *PDGFRB* were found to be significantly higher in sub-Cluster 3. The GSEA underscored functional disparities between the two sub-populations. In sub-Cluster 0, pathways associated with metabolic activity, such as oxidative phosphorylation and glycolysis, were prominent, suggesting a metabolically active phenotype. Additionally, this sub-cluster showed enrichment in the reactive oxygen species pathway and TNF-α signaling and NF-κB, highlighting potential roles in cellular stress response and inflammation. Intriguingly, sub-Cluster 3 displayed enrichment in pathways related to cell cycle and development, such as mitotic spindle and G2M checkpoint, which may reflect a proliferation-associated state. The presence of pathways like estrogen response early and epithelial-mesenchymal transition indicates a potential involvement in tissue remodeling and hormonal responses. KEGG pathways, including focal adhesion, ECM receptor interaction, vascular smooth muscle contraction, and regulation of actin cytoskeleton, suggested cellular motility, contractility, and lineage transition. These divergent pathways between sub-Cluster 0 and 3 suggest a different functional state of PCs, with sub-Cluster 0 displaying PC classic characteristics of metabolic and structure maintenance within the lung tissue, while sub-Cluster 3 aligns with stress response engagement, including cellular proliferation, movement, and lineage transition. A full list of enriched pathways identified in PCs from sub-Cluster 0 and sub-Cluster 3 can be seen in **Fig S9**.

Next, we re-analyzed idiopathic pulmonary arterial hypertension (IPAH) PCs and control PCs utilizing previously published scRNAseq data sets and analysis methods.^31,32^ Among the sub-reclustered 13 mural cell clusters, only sub-Clusters 5 and 6 had expressions of *HIGD1B* (**Fig 5C, left**). Voline plots show that sub-Cluster 5 is *HIGD1B ^low^ PDGFRB ^high^ as defined as* Type 2 PCs whereas sub-Cluster 6 is *HIGD1B ^high^ PDGFRB ^low^* as defined as Type 1 PCs. Peudotime analysis suggests that Type 2 PCs are lineage active by purple/green, whereas Type 1 PCs are quiescent and have mature lineage status by yellow using the color code scale (**Fig 5C, right**). Differentially expressed genes (DEG) to compare IPAH Type 2 PCs with healthy Type 2 PCs only revealed two significantly altered genes (adjusted *P<0.02*), which are *VIM* (**Fig 5D**) and *ITM2C*. IPAH Type 1 PCs compared to healthy Type 1 PCs also only show two significantly altered genes (adjusted *P<0.02*), which are *MT2A* and *MT1M*, When re-analyzed hypoxic murine mural cells scRNAseq datasets from our recent published work^31^, we identified PC clusters from other mural populations using high expression levels of *Higd1b* (**Fig 5E**). Due to limited cell numbers, we can not further sub-cluster the murine PC population. By DEG analysis, *Vim* was significantly upregulated in 3.4-fold hypoxic PCs than normoxic PCs(**Fig 5F**).

### Type 2 PCs express upregulated levels of Vimentin after 3wk Hx

We then performed confocal microscopy and IF staining in normoxic and hypoxic *Higd1b-tdT+/-* mice to determine if Vimentin would be upregulated in PCs. Under normoxic conditions, Type 2 PCs (**Fig 6A Panel a’’**) were close to a SMA+ arteriole and had a decreased number of processes compared to Type 1 PCs located in the parenchymal capillaries (**Panel a’**). Vimentin was seen scattered throughout the parenchyma. Under increased magnification, there was no expression of Vimentin in Type 1 or Type 2 PCs (**Fig 6A, Panel a’&a’’**). However, after exposure to Hx, Vimentin was upregulated in the out layer of distal arterioles and only in Type 2 PCs, which wrapped around a muscularized distal arteriole (< 50 µm) coexpressing SMA (**Fig 6A, panel b’’**). This staining result was consistent with the Volin plot results seen in **Fig 5D**. Type 2 PCs also demonstrated distinct morphology (**Panel b”**, spindle shape and shorter processes) compared to normoxic Type 2 PCs(**Panel a’’**, an oval body and thin processes). Hypoxic Type 1 PCs showed an increased number of processes (**Panel b’**) compared to normoxic Type 1 PCs (**Panel a’**). Taken together, our scRNA-seq analysis and experiments in PC lineage tracing mice using *Higd1b* as a specific PC marker suggested upon Hx, a subset of PCs (Type 2) relocated in the distal arterioles and express higher levels of Vimentin and lower levels of typical PCs genes including *Higd1b, Cspg4,* and *Pdgfrb.* This was in contrast to Type 1 PCs, which, under physiological conditions, have a low relative expression of *Vimentin* and higher expressions of *Higd1b, Cspg4,* and *Pdgfrb* (**Fig 6B**). These findings suggest, for the first time, a heterogeneity of PCs in the capillary networks and that treatments targeting PCs and/or mural cells may affect a wide range of cells and may lead to disruption of the microvasculature homeostasis and structure support rather than reversing the vascular remodeling seen in PH. Instead, therapeutic approaches should be considered to exclusively target Type 2 PCs and have no effects on Type 1 PCs.

**Figure 6:**
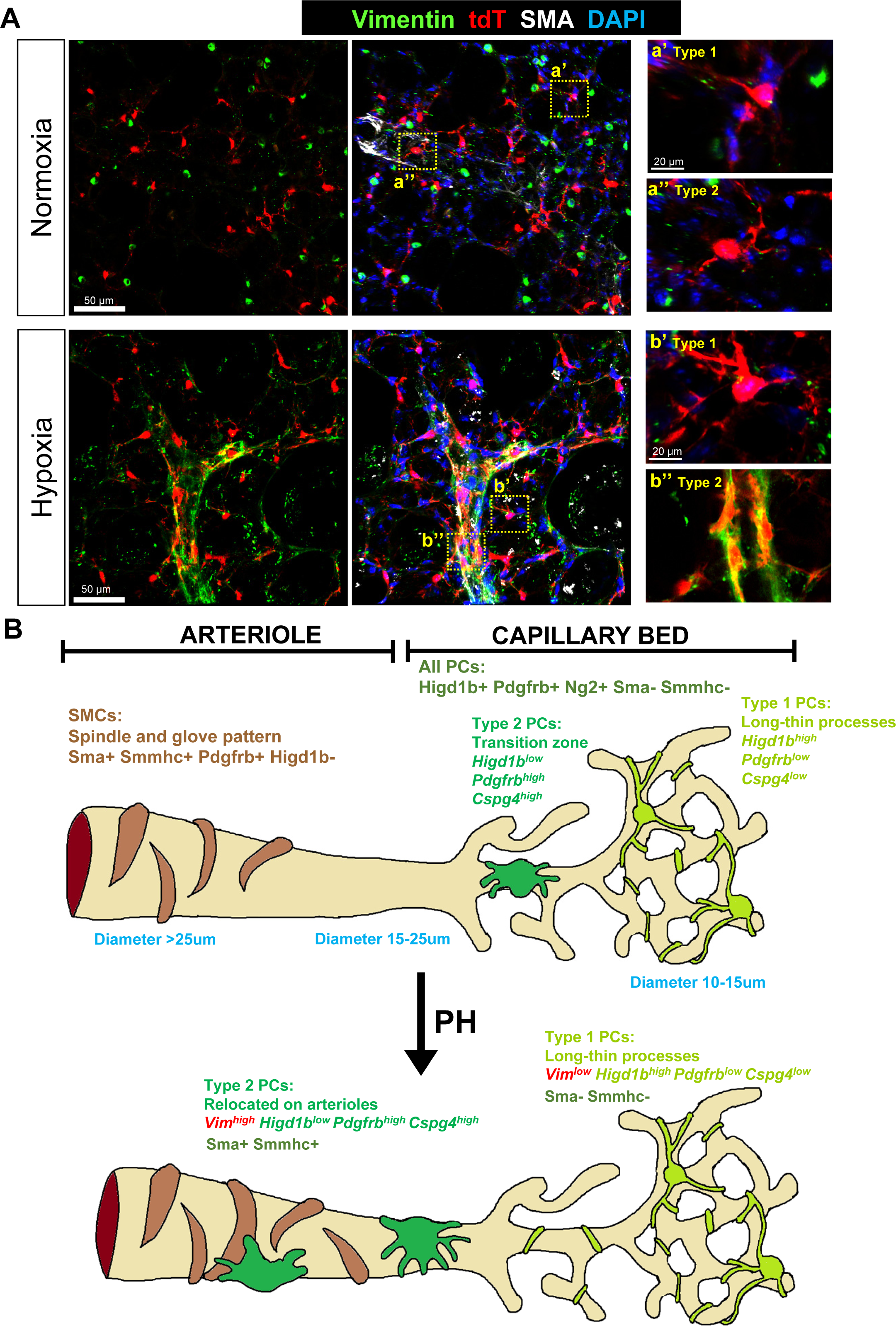
Type 2 PCs express upregulated Vimentin after Hx. (A) PCLSs were obtained from *Higd1b-tdT+/-* murine lungs exposed for 3wks of Hx or normoxia and stained for Vimentin (green), SMA(white) and nuclei (DAPI, blue). Scale bar: 50 µm. Panel a’ shows a Type 1 PC, with long and thin processes, which is located in the capillaries. Panel a’’ shows a Type 2 PC, which is located close to an SMA+ arteriole. Panel b’ shows a hypoxic Type 1 PC, with longer and thicker processes than normoxic Type 1 PC, located in the capillaries. Panel b’’ shows hypoxic Type 2 PCs, which accumulate and wrap around a SMA+ (white) arteriole, coexpressing Vimentin(green). Only hypoxic Type 2 PCs coexpress Vimentin. Scale bar: 20 µm. (B) The proposed model shows the location of PC subtypes and arteriolar SMCs in the pulmonary vasculature under physiological conditions and in the development of PH. Type 2 PCs upregulate vimentin in Hx-induced PH and have increased motility and lineage activity. Italic: gene expression. Non-Italic: protein expression.

## Discussion

In this study, we identified *Higd1b* as a unique and specific PC marker for both humans and mice in the cardiopulmonary system using scRNA-seq, spatial transcriptomics, and RNAscope (**Fig 1**). Utilizing the Cre-LoxP and CRISPR technologies, we generated a novel, tamoxifen inducible knockin mouse line (*Higd1b-CreERT2*), which effectively and specifically labeled PCs in abundance in the lung and heart (**Fig 2&3**). Through lineage tracing studies, we demonstrated two subtypes of PCs within the pulmonary capillary networks capable of dynamic transition (**Fig 4 &5**). Type 1 PCs located within the capillary network (diameter <10µm) maintain capillary homeostasis and integrity and were quiescent. Type 2 PCs located the zone between capillary and arterioles (diameter <25µm) and were lineage active and had multipotent properties, including the ability to transition into SMC-like cells, relocating to the arterioles in response to Hx, contributing to vascular remodeling and the development of Hx-induced PH (**Fig 6**).

PCs are an essential component of microvasculature and play key roles in regulating capillary homeostasis, angiogenesis, immune cell recruitment, vascular remodeling, and maintenance of the blood-brain barrier.^9,33–35^ They have been implicated in the development of numerous disease processes throughout the body, including PAH, Alzheimer’s disease, and diabetic retinopathy. A significant limitation and challenge in studying PCs is the lack of a clear and specific cell marker to distinguish them from other mural cell populations. Genetic mouse models, combining cell markers, fluorescent reporters, and lineage-tracing lines, represent powerful tools for genetically labeling PCs and tracking their behavior during development and in pathological conditions. Commonly used promoters for PC labeling include *Pdgfrβ*^36,37^*, Cspg4*^7,38,39^, *Tbx18*^39^, *LepR*^40^, Alkaline Phosphatase AP^40^, *Myh11*^41^, *Foxj1*^41^, and *Foxd1*^42^ and even dual reporter systems by *Pdgfrb-flp* and *Cspg4-FSF-CreER*^42^ or *PdgfraDreER* negative *PdgfrbCreER* positive^43^. However, it is essential to emphasize that none of these markers are exclusive to PCs, as SMCs, FBs, myofibroblasts, and other non-mural cells share these markers with PCs. Consequently, PC lineage-tracing studies cannot definitively rule out potential contamination from other cell types. scRNA-seq is an emerging and powerful tool for analyzing the genetic signature of individual cells, enabling the identification of genes for future diagnostic and therapeutic strategies. Through data mining from publicly available sources, we have identified *HIGD1B* as a unique and specific PC gene. Combining scRNA-seq with corresponding spatial transcriptomic data, our goal was to determine the subcellular location of mRNA molecules for *CSPG4, PDGFRB*, or *HIGD1B* and assign different cell types to their locations in a normal human lung section. *CSPG4* or *PDGFRB* was located around large-sized vessels, predominantly highlighted by Cluster 8, while *HIGD1B* was mostly found in the parenchyma and tentatively around capillaries (**Fig 1F**). The HIG1 hypoxia inducible domain (HIGD) gene family is comprised of five genes: *Higd1a, −1b, −1c, −2a, −2b.*^44^ *Higd1a* is regulated by hypoxia-inducible factor-1 (HIF1) and has roles promoting cell survival under hypoxic conditions.^45,46^ Previous studies have indicated that *Higd1b* promotes cardiomyocyte survival by maintaining mitochondrial integrity and its overexpression promotes cell survival through altered activation of caspase-3 and −9, but its role in pulmonary cells is currently unknown.^44^ Notably, PCs in different organs reside at different locations throughout the capillary microcirculation, thus leading to distinct morphological characteristics and signature genes.^47^ Our previous work suggested that PC markers may be organ specific and conserved in human and mouse hearts and lungs.^17^ The difference in PC gene expression between organs warrants further investigation in the future, as specific gene expression patterns in PCs may control cell dedifferentiation and be an important mechanism to understand when developing PC specific treatment strategies. Intriguingly, the *Higd1bCreERT2* mice truly reflect the PC specificity in different organs, as *Higd1bCre+* were abundant in hearts and lungs with decreased labeling in other organs, including the brain, retina, skeletal muscles, kidney, liver, and pancreas. Therefore, *Higd1b/HIGD1B* is an ideal cell marker to study PCs contribution to cardiopulmonary diseases.

Due to their close origin, PCs giving rise to SMCs or other mural cells have been extensively studied in many organs during development and under pathological conditions. For example, PCs coordinate the behavior of epithelial and vascular cells during lung morphogenesis.^48^ PCs serve as a source of SMC precursors during collateral artery formation in heart development.^38^ Resident PCs in postnatal skeletal muscle play a crucial role in contributing to differentiation in both the smooth muscle layer of blood vessels and the development of skeletal muscle fibers.^49^ FoxD1-lineage PC-like cells also contribute to the myofibroblast population following bleomycin-induced lung injury.^48^ PCs give rise to microglial cells after ischemic brain injury.^50^ PCs participate in vascular and fibrotic remodeling after ischemic damage in the heart.^51^ PCs in kidneys differentiate into myofibroblasts which contribute to collagen deposition and fibrosis.^14,35,52,53^ In contrast, some studies suggest that endogenous PCs in the heart, brain, skeletal muscle, and fat tissues do not behave as multipotent tissue-resident progenitors^54^. These discrepancies may be due to different PC subtypes, suggesting a potential diversity of plasticity and function. However, the classification of proposed PC subtypes remains challenging, again, due to the lack of PC-specific markers. Without a consensus, PCs have been defined by the inclusion and exclusion of numerous markers. PCs in the cerebral circulatory system closer to the arteriole end of the capillary bed may be involved in regulating blood flow, while PCs on the capillary bed more vital to maintaining the function of the blood-brain barrier, and PCs at the venule end of the capillary network regulate immune cell infiltration.^33,55–58^ Skeletal muscle Type 1 (Nestin-NG2+) PCs are fibrogenic and adipogenic in old and diseased muscle, while Type 2 PCs (Nestin+NG2+) generate new muscle tissue after injury.^59^ Additionally, Type 2 PCs recover blood flow in a mouse model of hindlimb ischemia.^60^ Two types of PCs specified as CD274+ capillary and DLK1+ arteriolar PCs are differentiated from human pluripotent stem cells.^61^ Applying scRNA-seq to IPAH and control lung cells, two subgroups of PCs were identified (**Fig 5C**). Type 1 are classical or synthetic PCs enriched with *HIGD1B* but with lower expression of SMC signature mRNAs, including *PDGFRB*, *ACTA2*, and *MYH11*. They have long punctate processes that partially wrap around capillary ECs within the basement membrane. They reside only on capillary beds, with one cell covering roughly 3-4 capillary nets. Type 1 PCs maintain capillary homeostasis and inhibit EC proliferation under normal conditions. They are fully differentiated and quiescent, thus having fewer multipotent properties. Under hypoxic conditions, their process surface area expands, and they wrap tighter around capillary EC junctions in response to arteriolar vasoconstriction, protecting against capillary leakage. Selective ablation of brain PCs provokes exuberant extension of processes from neighboring PCs to contact uncovered endothelium regions.^62^ We speculated that Type 1 PC process expansion may be a dynamic event similar to brain PCs due to the absence/movement of neighboring Type 2 PCs, but it requires further experimental evidence.

Type 2 are contractile PCs with relatively lower expression of *HIGD1B* and higher expression of SMC signature mRNAs, including *PDGFRb*, *ACTA2*, and *MYH11*, compared to Type 1 PCs. They reside at the border zone of capillary nets and arterioles (15-25μm). Their processes are shorter, making detachment and transmigration easier. Similar to Type 1 PC inhibition to EC growth, Type 2 PC’s main functions may prevent arteriolar SMCs from migrating to capillaries and maintain the homeostasis balance of the SMC-PC-capillary compartments. Under hypoxic conditions, Type 2 PCs translocate out of the border zone and reside on arterioles (diameter >25μm) and coexpress Sma+ Smmhc+, suggesting they have multipotent stem cell properties under pathological conditions and may be more motile SMC-like in response to arteriolar vasoconstriction. Vimentin orchestrates cytoskeletal and microtubule rearrangements and mechano-signaling to promote cell migration and polarity.^63–65^ Its upregulation in hypoxic Type 2 PCs may reduce cell-cell contact and drive cell translocation. In a recovery model where mice are re-exposed to normoxia after 3 weeks of Hx, PH is completely reversed. Intriguingly, the vast majority of Type 2 PCs are no longer required for remodeling and vasoconstriction and move back to the border zone, while Type 1 PC processes restore back to their normal size.

More studies are required to distinguish Type 1 and Type 2 PCs using unique markers and to further characterize and validate their function and plasticity. Their cellular and genetic identities should be further validated using clonal expansion and multiple color reporter systems, such as Confetti reporter lines. Additionally, subtypes of PCs in the capillary circulation of human lungs need to be carefully examined and characterized. Despite its advantages, scRNA-seq does not describe cell location and spatial information. Understanding PC spatial orientation is crucial for comprehending PC distribution or morphological changes in disease pathogenesis, especially if we can identify them in patient biopsy samples in relation to disease progression and severity. The other major limitation is that the geographic differences of human donor PCs regarding sex, age, and ethnicity are not explored. To more specifically label PC subtypes, barcoding cells with synthetic DNA sequences such as DARLIN, an inducible Cas9 barcoding mouse line, may be a useful tool to generate massive lineage barcodes across tissues and enable the detection of edited barcodes in profiled single cells.^66^ In future studies, the molecular mechanisms of Vimentin regulation in PC migration and polarity are necessary. PC lineage changes in other lung diseases such as chronic obstructive pulmonary disease (COPD) and idiopathic pulmonary fibrosis (IPF) should be carefully examined and investigated. Additionally, the role of PCs in the heart microvasculature is overlooked under pathological conditions. Targeting PC malfunction reveals enormous therapeutic possibilities, as PC-directed therapies have the potential to reverse or prevent disease progression and development in multiple scenarios. For example, human pluripotent stem cell-derived PCs have been successfully engrafted into the vasculature of ischemic murine limbs and promoted vascularization and muscle regeneration, highlighting the high ceiling of their potential as a therapeutic target.^67^ The proliferation of SMCs in PAH may be targeted using a PDGFRβ inhibitor (such as imatinib). However, PDGFRβ is expressed not only on SMCs but also on PCs, which play a crucial role in maintaining capillary homeostasis. Therefore, reducing the PC population with a PDGFRβ inhibitor may disrupt capillary homeostasis, leading to unexpected side effects and vascular leakage. To specifically target SMC proliferation, it may be more effective to use surface markers that are more specific to SMCs and exclude PC inhibition.

In summary, we identified *Higd1b* as a specific and unique cell marker able to differentiate PCs from other mural cell populations. We generated a novel, tamoxifen-inducible, PC reporter mouse *Higd1b-CreERT2* and confirmed the selective and appropriate labeling of PCs dominantly in the lungs and hearts. Our study, for the first time, suggests the existence of PC subtypes in the pulmonary circulation with different functions and dynamic changes in their morphologies under hypoxic conditions. This new cell marker and knockin mouse will provide the field of vascular biology with the necessary tools to perform fate mapping and gene knock-out experiments specific for PCs and lead to new discoveries on their contribution to disease development.

## Materials and Methods

### *HIGD1B* expression in Single-cell Databases Annotating PCs

We employed integrated datasets from the ‘Human Lung Cell Atlas (core)’ and the ‘Tabula Muris Senis’ to investigate the expression of *CSPG4*, *PDGFRB*, and *HIGD1B* in PCs of human and mouse lungs, respectively^22,68^. Additionally, single-cell sequencing data from Litvinukova et al. (2020) and Feng et al. (2022) were utilized for analogous investigations in human and mouse hearts^24,69^. These datasets, which included annotations for PCs, were derived from healthy subjects or mice. We used the Seurat R package to specifically examine the expression of *CSPG4*, *PDGFRB*, and *HIGD1B* across different cell types utilizing dot plots and density plots, employing the original UMAP coordinates from each dataset for visualization^70^. The analysis methods applied for Figure 5C-F were previously published in PMID: 38243138.

### Spatial Analysis of *HIGD1B* in lung tissue

We analyzed the ‘Non-diseased Lung’ dataset from the ‘Xenium Human Lung Preview Data’ to examine *CSPG4*, *PDGFRB*, and *HIGD1B* expression in lung tissue. The dataset underwent normalization with SCTransform, followed by dimensionality reduction using Principal Component Analysis (PCA)^70,71^. We selected the top 30 principal components based on the percentage of variance explained for subsequent analysis. These components were used to project the cells into a two-dimensional space using the Uniform Manifold Approximation and Projection (UMAP) algorithm^72^. Unsupervised clustering was performed with the FindClusters function (resolution set at 0.3), identifying distinct cell clusters^70^. We then utilized dot and density plots to investigate the expression patterns of *CSPG4*, *PDGFRB*, and *HIGD1B* across these clusters. For spatial visualization, a specific lung section (coordinates: x: 1200-2900, y: 3750-4550) was chosen to illustrate cell segmentation boundaries and the localization of individual *CSPG4*, *PDGFRB*, and *HIGD1B* molecules. Cells within this section were color-coded according to their cluster membership, enabling a detailed examination of spatial expression patterns.

### Sub-clustering Pericyte cluster utilizing Harmony

The Pericyte cluster, annotated from respiratory airway and lung parenchyma tissue was extracted from the ‘Human Lung Cell Atlas (core)’. We applied PCA to this focused dataset, selecting the top five principal components based on their contribution to variance. To mitigate the effects of potential confounders, Harmony was utilized, adjusting for variables including ‘dataset’, ‘assay’, ‘tissue sampling method’. ‘sequencing platform’, ‘development stage’, ‘tissue’, ‘subject type’, ‘study’, ‘lung condition’, ‘sex’, ‘self-reported ethnicity’ and ‘age or mean of age range’^30^. These principal components were subsequently used to project the cells into a two-dimensional space via the UMAP algorithm. Following this, unsupervised clustering was conducted using Seurat’s FindClusters function, with the resolution parameter set to 0.2. We then utilized dot and density plots to investigate the expression patterns of *CSPG4*, *PDGFRB*, and *HIGD1B* across these Pericyte sub-clusters.

### Differential Gene (DE) Expression Analysis and Gene Set Enrichment Analysis (GSEA)

DE analysis was carried out to compare Pericyte sub-clusters 0 and 3 using the Wilcoxon rank sum test^70^. In this analysis, a positive log2 fold change (log2FC) indicates higher gene expression in sub-cluster 0 relative to sub-cluster 3. DE genes were ranked in descending order based on their log2FC to create a pre-ranked gene list for subsequent analysis. We employed the GSEApy to perform GSEA on this pre-ranked gene list against curated gene sets from the Hallmark collection, and the Kyoto Encyclopedia of Genes and Genomes (KEGG)^73^. Statistical significance for pathway enrichment was assessed using a permutation test, with pathways exhibiting an adjusted q-value below 0.05 considered statically significant.

### Data Availability

For experiments pertaining to scRNA-seq, microfluid, droplet based data was processed using the 10× Genomics platform (10×3′v2) of the (1) Tabula Muris Senis, (2) Human Lung Cell Atlas, (3) Adult Human Heart, and (4) The Mouse Heart datasets, which were obtained from the Cellxgene collections (https://cellxgene.cziscience.com) (<u>16</u>, <u>24</u>, <u>25</u>). The pre-processed, publically available scRNA-seq datasets utilized in experiments include:

1. Tabula Muris Senis compendium for murine lung tissue: (https://cellxgene.cziscience.com/collections/0b9d8a04-bb9d-44da-aa27-705bb65b54eb).
2. The Human Lung Cell Atlas dataset for human lung tissue: (https://cellxgene.cziscience.com/collections/5d445965-6f1a-4b68-ba3a-b8f765155d3a).
3. The Adult Human Heart dataset for human cardiac cells: (https://cellxgene.cziscience.com/collections/b52eb423-5d0d-4645-b217-e1c6d38b2e72).
4. The Mouse Heart dataset for murine cardiac cells: https://www.ncbi.nlm.nih.gov/geo/query/acc.cgi?acc=GSE193346.

Spatial transcriptomics of the human lung was obtained from the Xenium Human Lung Preview Data (Non-diseased Lung) using 10x Genomics Datasets which can be found at: (https://www.10xgenomics.com/datasets/xenium-human-lung-preview-data-1-standard).

### Experimental animals

All animal procedures conducted in this study were approved by the Institutional Animal Care and Use Committee (IACUC) guidelines at Boston Children’s Hospital (BCH) and adhered to the published guidelines of the National Institutes of Health (NIH) on the use of laboratory animals. Mice were housed in the animal facility at BCH with a 12-hour light/dark cycle and ad libitum access to rodent chow and water.

The founder #25 *Higd1b-CreERT2* was crossbred with Ai14 (*Higd1b-CreER::R26-tdTomato)* for generation of the *Higd1b-tdT* and also crossbred with the *Rosa26-mTmG*(https://www.jax.org/strain/007576) to generate *Higd1b-mTmG*. Ear clip-based genotyping was used to identify knockin and control mice. *Cspg4-CreERTM*(https://www.jax.org/strain/008538) was crossbred with Ai14 for the generation of the *NG2-tdT* and tamoxifen induction following the previously published protocol.^7^

### Donor plasmid and Guide RNA

Higdb1-P2A-CRE-ERT2 targeting vector (donor) was generated by assembling 1kb left (LHA) and 1kb right homology (RHA) arms flaked by P2A-CRE-ERT2 (Fig 1-A) and designed and purchased by Vectorbuilder Inc(cat# VB220531-1072sdb). Two gRNA (27F & 37F) were used for CRISPR/Cas9 mediated knock-in into Higdb1 exon 4. 0.61pmol each of crRNA and tracrRNA was conjugated and incubated with Cas9 protein (30ng/ul) to prepare ribonucleoprotein complex (RNP) according to Aida et al and donor DNA (10ng/ul) was mixed with RNP for microinjection cocktail.

### Microinjection

For knock-in mouse generation - microinjection cocktail was injected into 0.5dpc embryos harvested after mating C57Bl6-Hsd (Envigo). Post-injection embryos were reimplanted into CD1 (Envigo) pseudo-pregnant foster females and allowed to term. Tail snip biopsies were collected from pups at P7.

### Genotyping

Tail snip genomic DNA was prepared from pups and analyzed by PCR using the following primer pairs (Fig 2C). Primer pairs LF + C1 for LHA, C2 + C3 for Cre, C4 + RR for RHA. PCR reaction containing 0.5uM each primer pair, 1xQ5 MM buffer (NEB), 100ng genomic DNA and was amplified using a thermocycler (BioRad): 95°C-3 min. 95°C −30sec. annealing 67 °C to 72 °C, 72 °C 1 min. for 35 cycles and final extension at 72 °C for 5 min. PCR products were analyzed on 1% agarose gel (Seakem - GTG), 1XTAE buffer, and DNA was purified from gel using a Gel Extraction kit (QIAQuick - Qiagen) and cloned using TOPO cloning Kit (Invitrogen).

Plasmid DNA was prepared using miniprep kit (Qiagen) and Sanger sequenced using primers: S1 forward primer outside of LHA and S2 reverse primer at the 5’ of CRE; S3 forward primer at the 3’ of CRE, S4 forward primer in ERT2 and S5 reverse primer outside of RHA.

### Reporter gene activation via tamoxifen injection

For tamoxifen induction, *Higd1b-tdT* mice were injected with 6 mg of tamoxifen dissolved in corn oil (20 mg/ml) over 3 days (143mg/kg for 30g mouse). The median age and weight of *Higd1b-tdT* male mice used in this study was 7.47 +/-1.22weeks and the average body weight 28.14 +/-1.17grams at the time of injection. The median age of female mice was 7.35+/-1.22 weeks and the average body weight 28.1+/-1.17grams.

*Higd1b-mTmG* mice were injected with 4 mg of tamoxifen dissolved in corn oil (20 mg/ml) over 2 days (∼133 mg/kg for 30g mouse). The median age and weight of *Higd1b-mTmG* male mice used in this study was 7.55 +/-1.04 weeks and the average body weight 28.53 +/-1.2 grams. The median age of female mice was 7.86 +/-1.04 weeks and the average body weight 28.55 +/-1.2grams at the time of injection.

All mice were allowed to rest for 7-14 days before being exposed to Hx or undergoing tissue harvest. As negative controls, tissue from mice (*Higd1b-tdT+/-* and *Higd1b-mTmG+/-*) without tamoxifen and wildtype mice (Cre positive flox negative or Cre negative flox positive as *WT-tdT+/-* and *WT-mTmG+/-*) injected with similar doses of tamoxifen were harvested and underwent inspection for endogenous PC labeling. Experimental animals are defined as heterozygous of Cre and flox.

### Hypoxia Studies

Mice were placed in a Hx chamber and exposed to 10% FiO_2_ with ad libitum access to rodent chow and water for up to three weeks. The environment within the chamber was established through a continuous mixture of room air and nitrogen gas. The chamber environment was continuously monitored using an oxygen analyzer (Servomex, Sugar Land, TX). CO_2_ was removed with lime granules, and the temperature was maintained between 22-24°C. The chamber was inspected at least daily for animal welfare, O_2_ concentration, CO_2_ concentration, and humidity.

### Vibratome tissue preparation

Animals were euthanized with controlled isoflurane and cervical dislocation. After euthanasia, mice were secured in the supine position, and a midline incision was made to expose the abdominal and thoracic cavities. The sternum was dissected to expose the contents of the mediastinum and then the abdominal aorta was located and severed. A 25G butterfly needle was inserted into the right ventricle (RV) and slowly perfused with 15cc of ice-cold 1X phosphate buffered saline (PBS) to flush the red blood cells from the circulatory system. Once the lungs were white in appearance, the trachea was cannulated and the lungs inflated with 2% low-melting point agarose in 1X PBS. After complete inflation, the trachea was tied and cold 1x PBS poured over the lungs to solidify the agarose and preserve the structure of the lung. The lungs were then carefully removed from the mediastinum and placed in 4% paraformaldehyde (PFA) at 4°C overnight, followed by washing in 1X PBS the next day. Lung lobes were then separated and sectioned with a vibratome machine (Leica VT1000 S) at a thickness of 300µm for IF staining and microscopy.^28^

The heart was removed from the mediastinum after flushing with 15cc of 1X PBS and the right and left atrium dissected to expose ventricles. Heart samples were washed in 1X PBS and then fixed in 4% PFA overnight at 4°C. After three washes in 1X PBS (one hour each), the samples were placed in 4°C overnight on a rotating plate. The next day, samples were removed and sectioned with a vibratome machine at a thickness of 300µm.

### Optical cutting temperature (OCT) tissue preparation

After appropriate euthanasia, the hind leg of the mice was secured and an incision was made into the subcutaneous tissue located over the femur to expose the muscle. Skeletal muscle from the femur, identified by its striations and orientation of muscle fibers, was carefully dissected and removed. For connective tissue preparation, the tissue surrounding the descending aorta in the abdominal cavity was identified and dissected after the removal of the mediastinal contents as previously described.^31^ Connective tissue and skeletal muscle were placed in 4% PFA at 4°C overnight and then washed in 1X PBS for another day.

Brain tissue was harvested from mice by first securing the mouse in the prone position after euthanasia and dissecting away the hair and subcutaneous tissue to expose the skull. The skull was carefully pierced at the estimated location of the sagittal suture, and the cranial bones were dissected, being careful not to damage the brain tissue underneath. Once an opening was created and the brain removed from the skull, the tissue was placed in 4% PFA at 4°C overnight. Tissues were washed in 1X PBS for an additional day.

The liver and kidney were located and carefully removed after euthanasia and flushing of red blood cells from the vasculature. The tissue was fixed in 4% PFA at 4°C overnight and then washed in 1X PBS.

All the above samples were completely submerged in a 30% sucrose/PBS solution and placed at 4°C overnight on a rotating plate for several days until they sunk to the bottom. They were embedded with 100% OCT and stored at −80°C for future experiments. OCT-prepared samples were then sectioned with a Cryostat machine (RWD, FS800A Cryostats) at a thickness of 10 um and mounted on microscopy slides for IF staining.

The heart was removed from the mediastinum after flushing with 15cc of 1X PBS and the right and left atrium dissected to expose ventricles. Heart samples were embeded in 100% OCT solution. The next day, samples were removed and sectioned with a Cyrostat machine at a thickness of 10µm.

Retina were dissected from mice after euthanasia and fixed in 4% PFA for one hour and then washed with 1x PBS overnight at 4°C. The samples were carefully dissected under a dissecting microscope to expose the optic nerve and surrounding vasculature. Fixed retina were stored at 4°C until IF staining. After staining, the samples were placed on microscopy slides with Prolong Gold Antifade Solution containing DAPI.

### Immunofluorescence staining

Vibratome-prepared precision cut lung sections (PCLSs) and heart tissues were blocked with 5% goat serum in 0.5% Triton X-100/PBS (PBS-T) for one hour at room temperature. Samples were incubated with primary antibodies diluted in 5% goat or donkey serum and 0.5% PBS-T at 4°C for 48 hours. PCLSs were then washed three times in 1X PBS (fifteen minutes per wash) followed by incubation with 488, 555, or 647 fluorophores containing secondary antibodies (1:250 concentration) overnight at 4°C. Samples were then washed again and placed on microscopy slides with Prolong Gold Antifade Solution containing DAPI (Life Technologies Corporation). After thawing to room temperature, OCT-prepared tissue and retina were stained following a similar technique. All images were captured using a Zeiss confocal 880 Airyscan 2 microscope and processed by Aivia software.

The following antibodies were used for IF staining:

Mouse-anti-mouse/human SMA-647 (1:100; sc-32251-AF647, Santa Cruz Biotechnology)

Rat-anti-mouse CD31(1:100; 553370, BD-Pharmingen)

Rabbit-anti-mouse PDGFR-β (1:100; MA5-15143, Thermo Scientific)

Rabbit-anti-mouse MYH11 (1:100; AB53219, ABCAM)

Isolectin GS-IB_4_-488 (1:100; I21411, Thermo Fisher Scientific/Invitrogen)

Goat-anti-mouse Cardiac Troponin 1 (1:100); ab56357, Abcam)

## Acknowledgements

We thank the IDDRC Gene Manipulation Core at Boston Children’s Hospital, funded by NIH P50 HD105351. We thank We thank the Neurobiology Department and the Neurobiology Imaging Facility for consultation and instrument availability that supported this work by NINDS P30 Core Center Grant #NS072030 from Harvard NIF core for imaging or analysis. We thank the Harvard Digestive Disease Center and NIH grant P30DK034854 in all publications and presentations emerging from HDDC core services, resources, technology, and expertise. We thank Joint Biology Consortium (JBC) of BWH/BCH grant number NIH/NIAMS 2P30AR070253 to support our work. NIH R01 HL141851, R01 HL145676 (to JCW); NINDS P30 Core Center Grant #NS072030 from Harvard NIF core for imaging or analysis; NIH/NIAMS 2P30AR070253 from JBC consortium; the Harvard Digestive Disease Center and NIH grant P30DK034854 in all publications and presentations emerging from HDDC core services, resources, technology, and expertise.

## Author Contributions

Conceptualization: KY; data curation: TK, YK, SHB, MB, YL, TL, KY; investigation and methodology: TK, YK, SHB, MB, YL, TL, KY; supervision: KY; writing-original draft: TK, SHB, MB, YK; providing reagents: JQ, JCW, BAR, VdJP; writing-review and editing: TK, YK, SHB, KY. All authors reviewed and approved the final manuscript.

## Supplemental Figure Legends

**Supplemental Figure 1:**
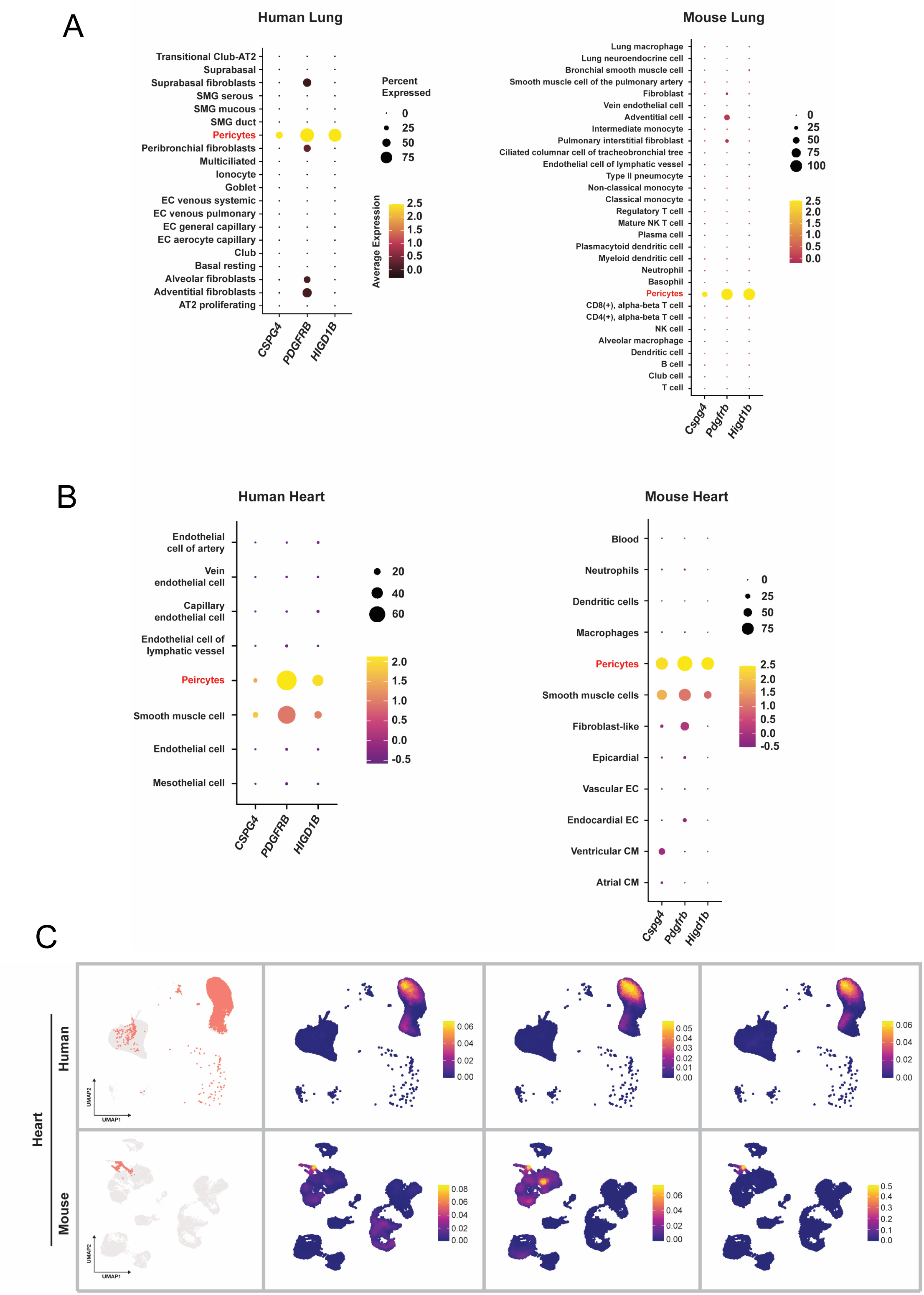
Human and murine lung scRNA-seq analysis demonstrates unique expression of *HIGD1B* to PCs. (A) Dot plots show expression levels of *CSPG4*, *PDGFRB*, and *HIGD1B* within all annotated cell types in human and mouse lung tissues. Note the expression of *HIGD1B* is exclusive to human and mouse PCs. (B) Dot plots show expression levels of *CSPG4*, *PDGFRB*, and *HIGD1B* within all annotated cell types in human and mouse heart tissues. (C) UMAP visualization of PC distributions within human and mouse heart tissues, using original UMAP coordinates and cell type annotations of single-cell sequencing data from Litvinukova et al. (2020) and Feng et al. (2022). The density plot specifically highlights the expression of *CSPG4*, *PDGFRB*, and *HIGD1B*.

**Supplemental Figure 2:**
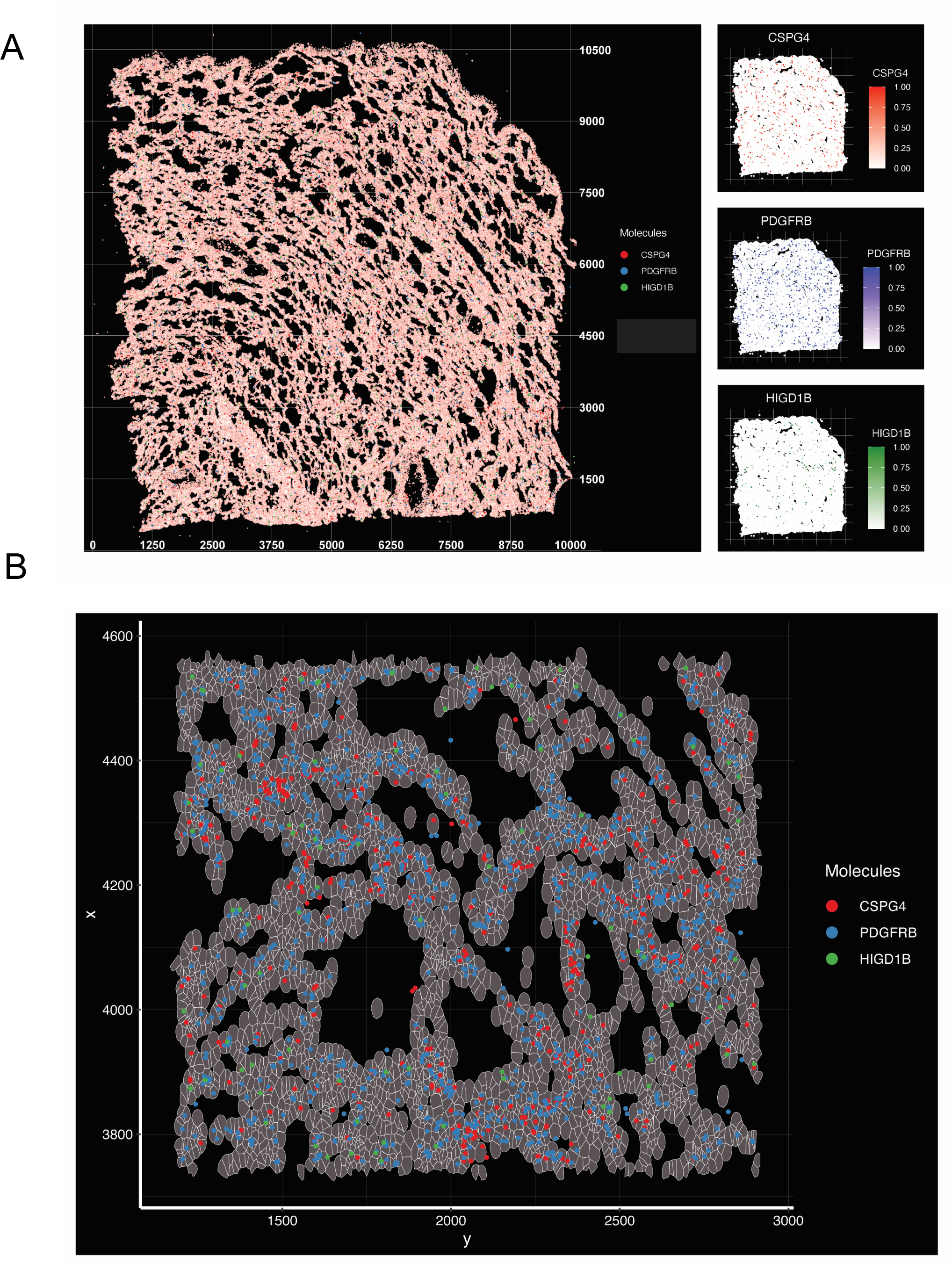
Spatial Analysis of PC markers in Non-Diseased Lung Tissue. (A) Spatial transcriptomic analysis shows the distribution of *CSPG4*, *PDGFRB*, and *HIGD1B* across the full lung tissue section.| (B) Magnified area of spatial transcriptomic map shows the distribution of *CSPG4(red)*, *PDGFRB (blue)*, and *HIGD1B (green)* across the selected lung tissue section (coordinates: x: 1200-2900, y: 3750-4550).

**Supplemental Figure 3:**
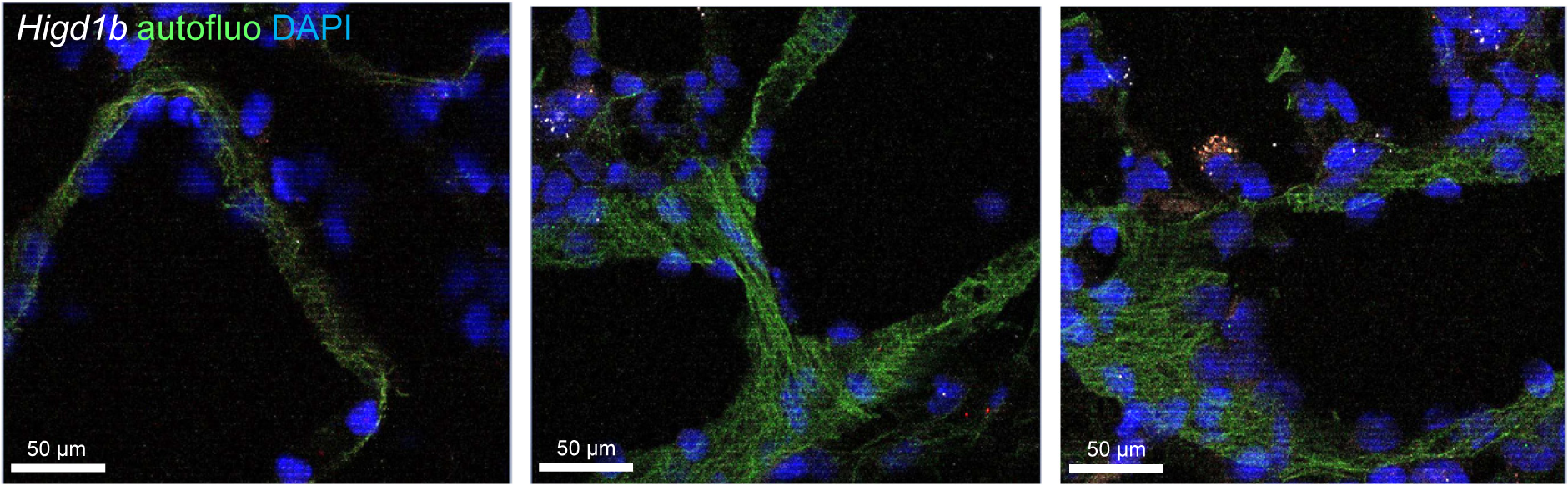
*Higd1b* mRNA expression is absent in arterial SMC layers. RNAscope shows the absence of *Higd1b* (white) expression in arterial SMC layers. Autofluorescence: green, indicating a SMC layer. DAPI: blue. Scale bar: 50 µm.

**Supplemental Figure 4:**
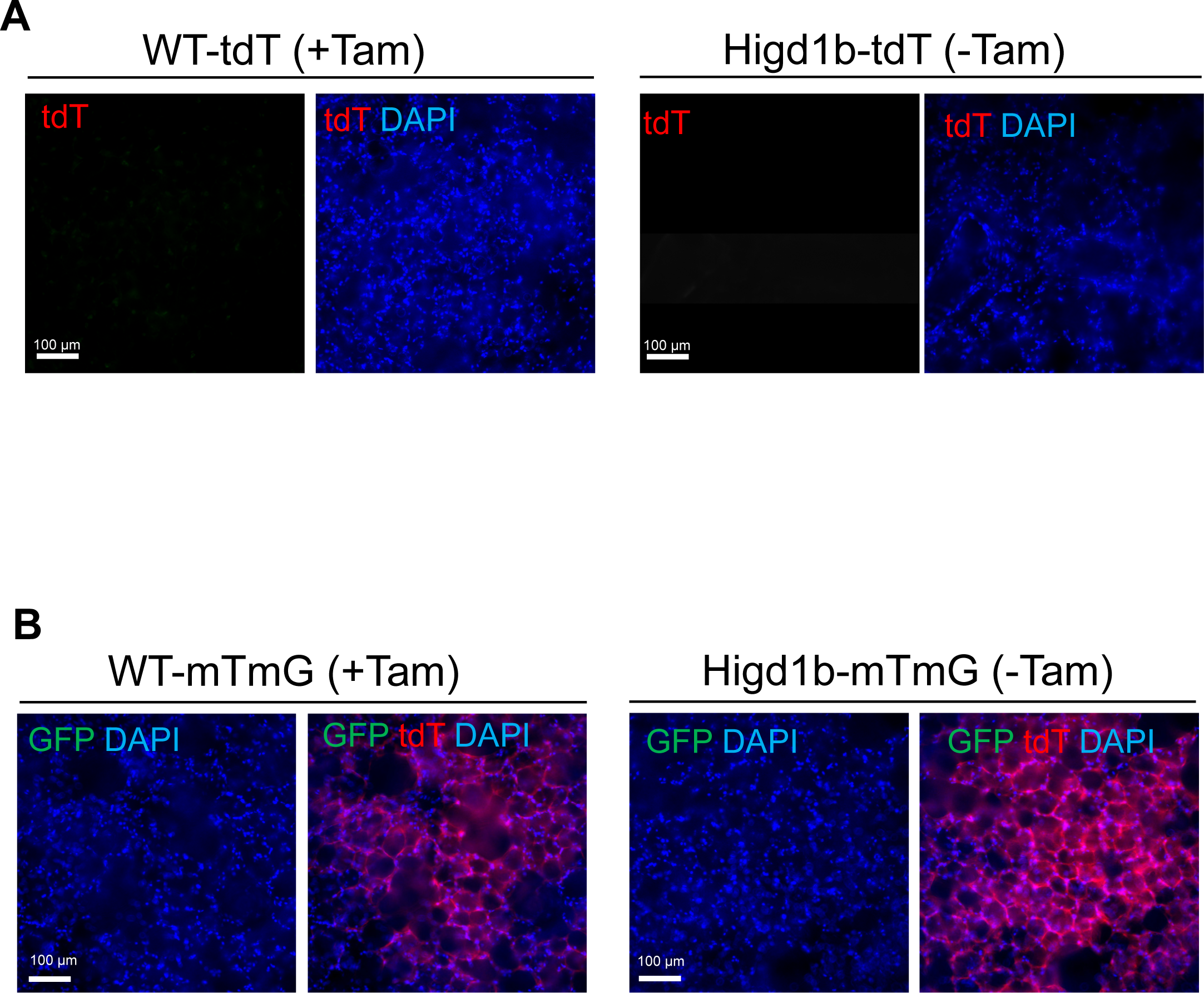
Control experiments for *Higd1b-tdT* or *Higd1b-mTmG* mouse lungs and WT lungs. (A) PCLSs from *WT-tdT+/-* mice after tamoxifen administration and *Higd1b-tdT+/-* mice without tamoxifen revealing an absence of tdT+ cells (red). DAPI: blue. Scale bar: 100 µm. (B) PCLSs from *WT-mTmG+/-* mice with tamoxifen and *Higd1b-mTmG+/-* mice without tamoxifen showing no membrane GFP reporter (green) expression but the presence of membrane tdT color(red). tdT reporter: red. DAPI: blue. Scale bar: 100 µm.

**Supplemental Figure 5:**
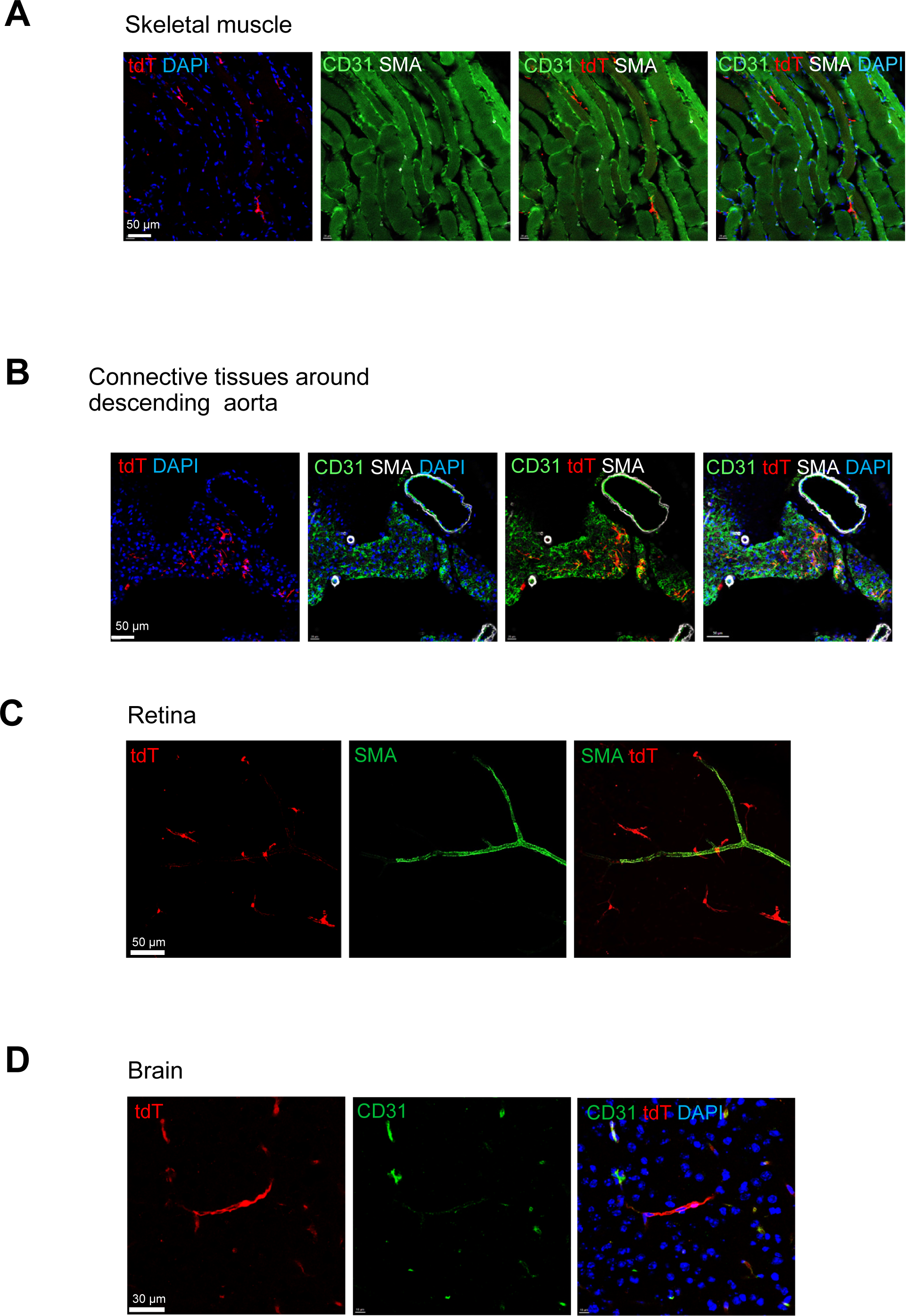
*Higd1b-tdT+/-* labels some PCs in other organs *in vivo*. (A) Skeletal muscle from *Higd1b-tdT+/-* mice with staining for CD31 (green), SMA (white), and DAPI (blue). tdT reporter: red. Scale bar: 50 µm. (B) Connective tissues surrounding the descending aorta from *Higd1b-tdT+/-* mice were stained with CD31: green, SMA: white, and DAPI: blue. tdT reporter: red. Scale bar: 50 µm. (C) Retina from *Higd1b-tdT+/-* mice and IF staining for SMA (green). tdT reporter: red. Scale bar: 50 µm. (D) IF staining of cerebral tissue from *Higd1b-tdT+/-* mice for CD31 (green) and DAPI (blue). tdT reporter: red. Scale bar: 30 µm.

**Supplemental Figure 6:**
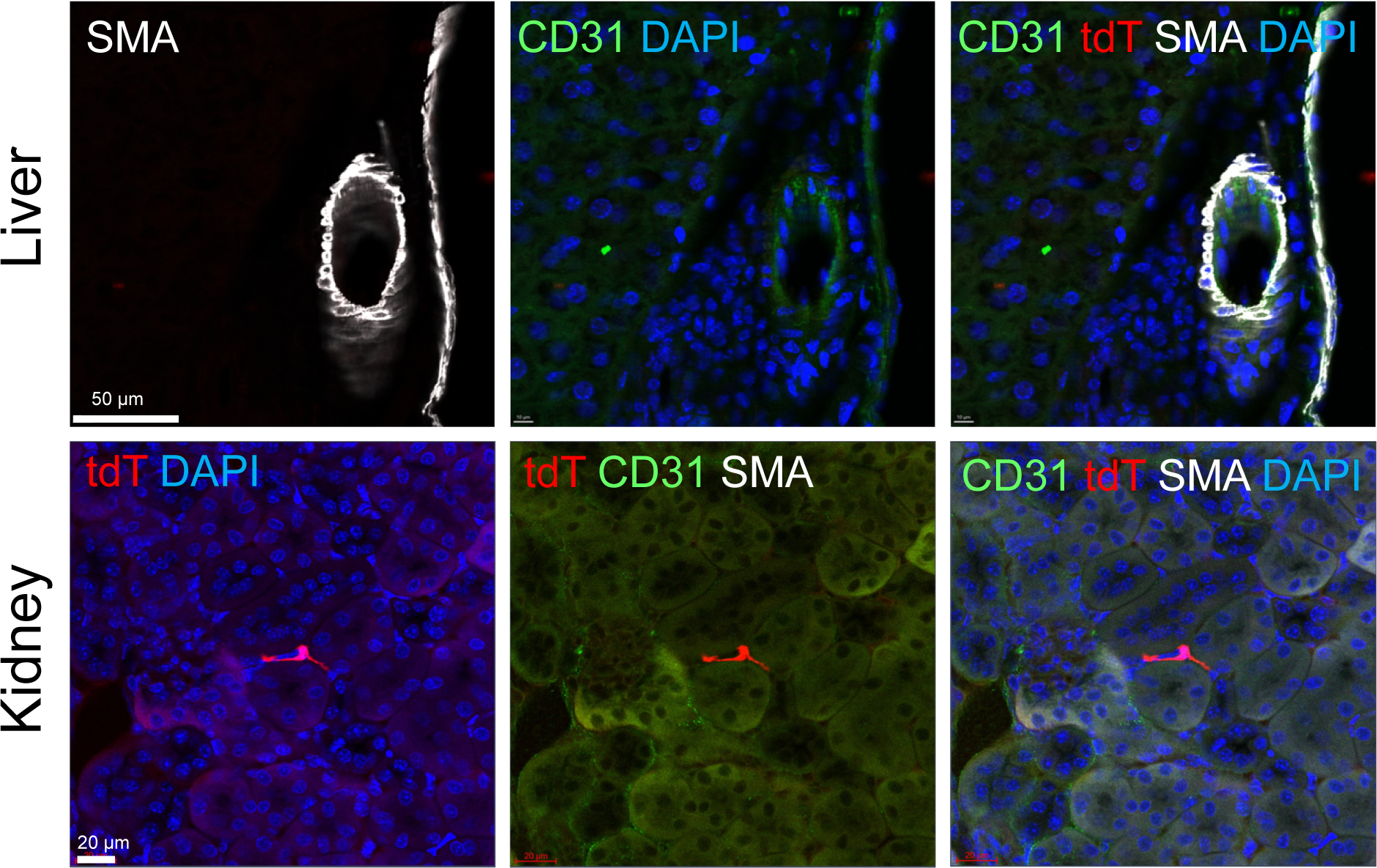
*Higd1b-tdT+/-* PCs are absent in the liver and the kidney. (A) Liver (top panel) and kidney (bottom panel) tissue from *Higd1b-tdT+/-* mice with staining for SMA (SMCs: white), CD31 (ECs: green), and DAPI (nuclei: blue) reveals a lack of tdT-positive PCs (red). Scale bar: 50 and 20 µm.

**Supplemental Figure 7:**
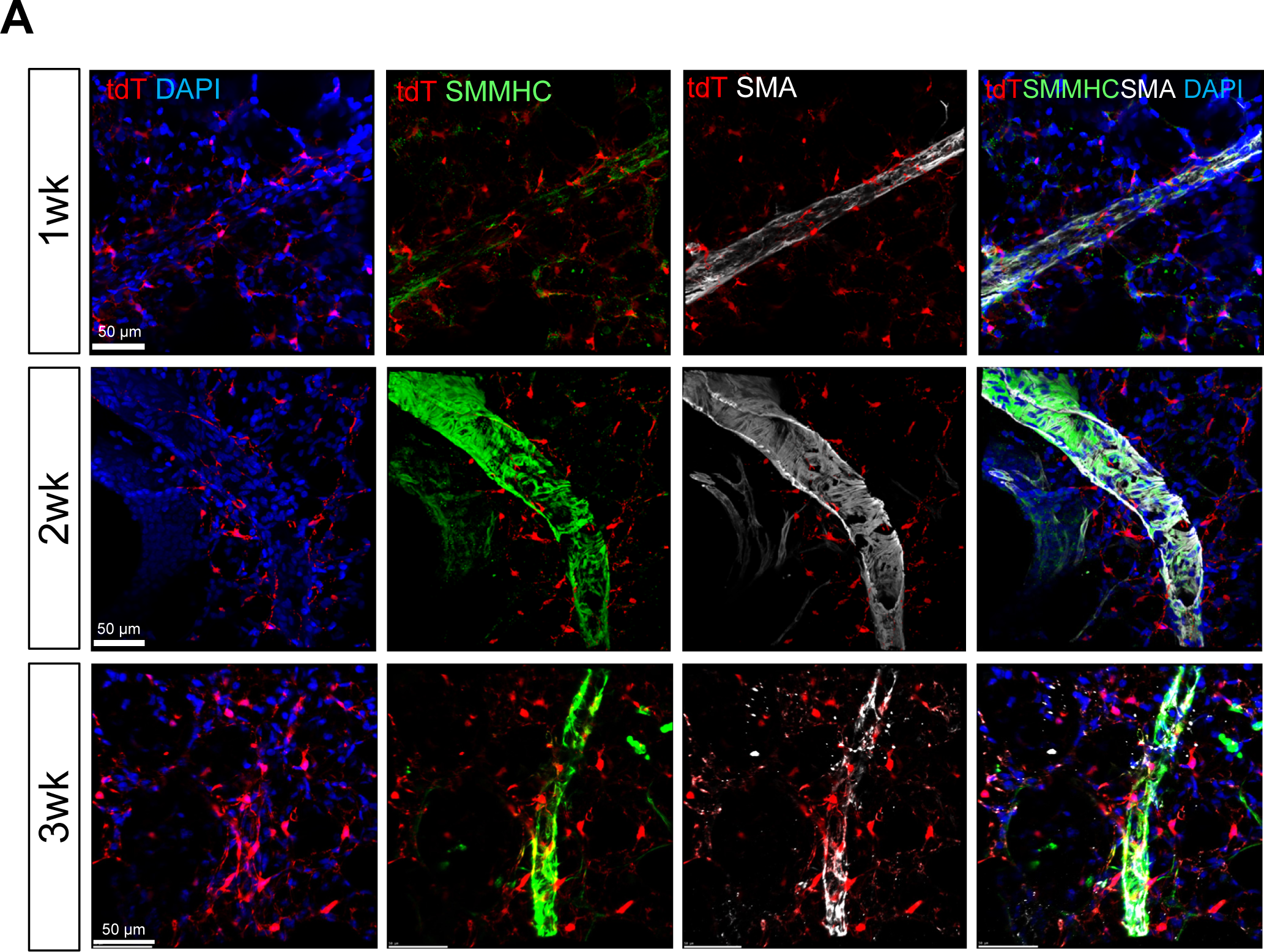
Lineage tracing shows that *Higd1b-Cre+* cells are accumulated in muscularized distal arterioles by different hypoxic exposure times. (A) Accumulation of tdT-positive PCs (red) in muscularized distal arterioles with staining for SMC markers SMMHC (green) and SMA (white) after 1, 2, and 3 weeks of Hx. DAPI: blue. Scale bar: 50 µm.

**Supplementary Figure 8:**
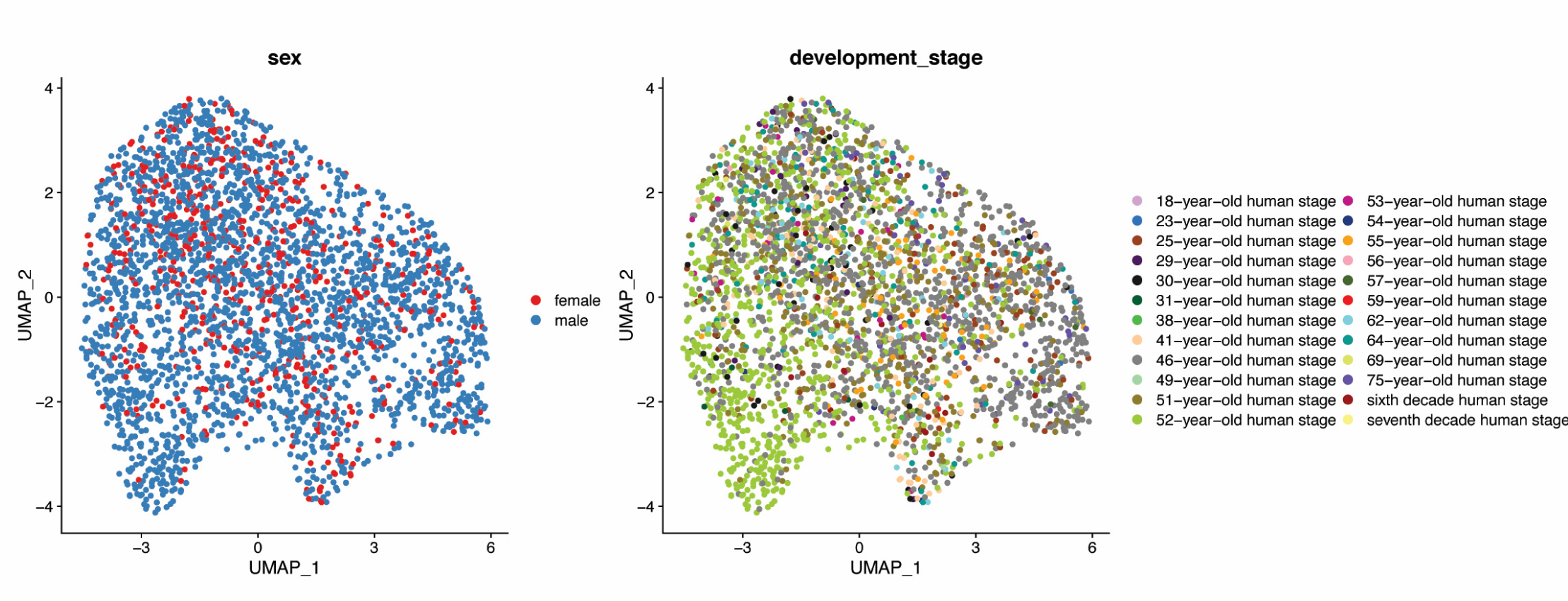
Patient demographics from annotated PC subclusters are included. UMAP visualization of Pericyte sub-populations within the annotated PC cluster from the ‘Human Lung Cell Atlas (core)’. The representation highlights the demographic distributions, categorizing cells according to the sex and age of the donors.

**Supplementary Figure 9:**
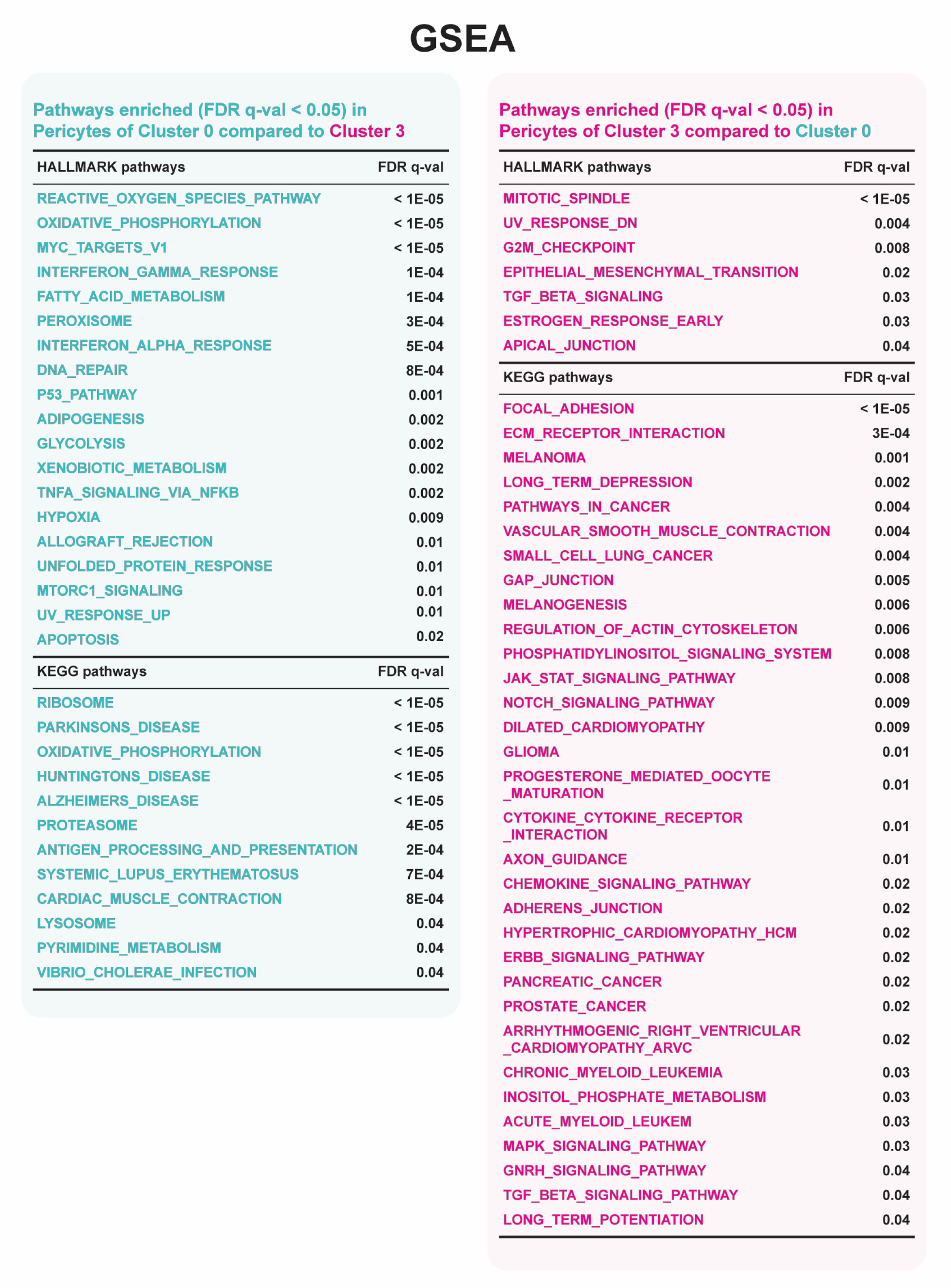
Gene Set Enrichment Analysis in PC sub-Cluster 0 and 3 is performed. A full list shows the results of Gene Set Enrichment Analysis (GSEA) comparing PC sub-Cluster 0 and 3 with an FDR below 0.05.

